# An island “endemic” born out of hybridization between introduced lineages

**DOI:** 10.1101/2022.09.14.507826

**Authors:** Jessie F. Salter, Robb T. Brumfield, Brant C. Faircloth

## Abstract

Humans have profoundly impacted the distribution of plant and animal species over thousands of years. The most direct example of these effects is human-mediated movement of individuals, either through translocation of individuals within their range or the introduction of species to new habitats. While human involvement may be suspected in species with obvious range disjunctions, it can be difficult to detect natural versus human-mediated dispersal events for populations at the edge of a species’ range, and this uncertainty muddles how we understand the evolutionary history of populations and broad biogeographic patterns. Studies combining genetic data with archeological, linguistic, and historical evidence have confirmed prehistoric examples of human-mediated dispersal; however, it is unclear whether these methods can disentangle recent dispersal events, such as species translocated by European colonizers during the past 500 years. We use genomic DNA from historical specimens and historical records to evaluate three hypotheses regarding the timing and origin of Northern Bobwhites (*Colinus virginianus*) in Cuba, whose status as an endemic or introduced population has long been debated. We discovered that bobwhites from southern Mexico arrived in Cuba between the 12th and 16th centuries, followed by the subsequent introduction of bobwhites from the southeastern USA to Cuba between the 18th and 20th centuries. These dates suggest the introduction of bobwhites to Cuba was human-mediated and concomitant with Spanish colonial shipping routes between Veracruz, Mexico and Havana, Cuba during this period. Our results identify endemic Cuban bobwhites as a genetically distinct population born of hybridization between divergent, introduced lineages.

## Introduction

Translocation of plants and animals has been a pervasive practice throughout human history (1), and genetic studies of the movement of domesticated and human-associated species have provided insights into population genetics (2), invasion biology (3), phenotypic evolution (4), and disease ecology (5). When combined with archeological data, translocations have also shaped our understanding of human history. For example, genetic data from moth skinks (*Lipinia noctua*) stowed away in prehistoric canoes provided strong support for the “express train” hypothesis of human colonization in the Polynesian islands (6). Similarly, genetic data from modern and historical herbarium specimens of sweet potatoes (*Ipomoea batata*) supported the archeological and linguistic evidence of pre-Columbian contact between Polynesians and South Americans (7).

Evidence of human translocations is commonly detected on islands due to their unique biotic assemblages and because water is often a significant barrier to terrestrial dispersal (8). For example, combined fossil, genetic, and archeological evidence suggested that northern common cuscuses (*Phalanger orientalis*) were introduced to several islands from New Guinea by humans ~20 kya, and other marsupial distributions on islands throughout Australasia may have been shaped by similar prehistoric human translocations (9, 10). For some species, the effects of human intervention on their distributions and demographic histories remain unclear, particularly when natural and human-mediated dispersal events occur within relatively recent timescales (e.g., the Holocene, 11.7 kya - present) and near the edges of a species’ native range (11, 12). For example, these factors have limited our understanding of how Cuban tree frogs (*Ostepilus septentrionalis*) reached Florida: a short time frame for natural dispersal (<11.7 kya) combined with 20th century human-mediated introductions have confounded our understanding of whether they naturally colonized the state (13). Studying this latter class of natural versus human-mediated dispersal events across recent (11.7 kya - present) timescales remains challenging, particularly when supporting fossils, genetic data, archaeological evidence, or historical records are few.

Here, we combine historical literature and genomic DNA from historical specimens to investigate an island population of uncertain origin – the Cuban population of Northern Bobwhites (*Colinus virginianus;* hereafter bobwhites). Bobwhites are small, sedentary quails native to southeastern Canada, the eastern United States, Mexico, Guatemala, and Cuba (Figure 1). Along with Cuba, bobwhites have been documented on at least eight other islands in the Greater Antilles (14), and, except for Cuba, there is broad consensus that these populations were introduced by Europeans during the last 500 years due to the low dispersal capabilities of bobwhites (15, 16) and the fact that bobwhites were not observed on these islands until relatively recently (17–19).

**Figure 1.**
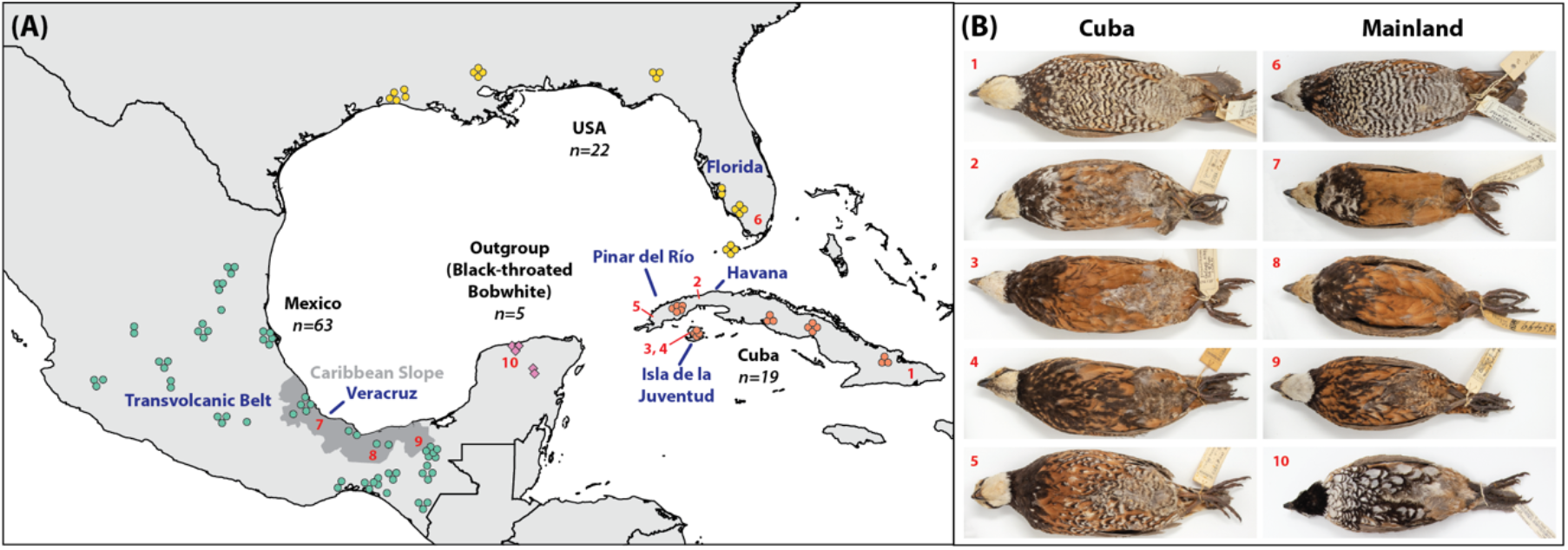
Sampling localities and example plumages of Northern Bobwhites and Black- throated Bobwhites. **(A)** Sampling localities and sample number per locality for 104 Northern Bobwhites and 5 Black-throated Bobwhites. Markers are colored by geographic population: USA (yellow); Mexico (teal); Cuba (orange); and Black-throated Bobwhites (outgroup; pink). Notable localities and geographic features mentioned throughout the text are labeled in blue; the Caribbean Slope of southern Mexico (including parts of Veracruz and Tabasco states) is shown in dark gray. Red numbers show collection localities of specimens in (B). **(B)** Example plumages of Northern Bobwhites (1–9) and Black-throated Bobwhites (10). Specimens 1-4 from Cuba have been paired with specimens 6-9 from mainland populations with similar plumages. Specimen 5 displays the unique *C. v. cubanensis* plumage described by Gould (21). Specimen photographs were taken at the National Museum of Natural History and are not represented in the genetic dataset.

Bobwhites were first observed on mainland Cuba and Isla de la Juventud (an island 50 km south of Pinar del Río and Havana provinces; Figure 1) by Europeans in 1839, and they were initially assumed to be introduced from populations in the eastern USA (20). Soon after, however, Cuban bobwhites were described as a distinct species, *C. cubanensis*, due to their smaller bills and their unique plumage pattern compared to bobwhites from mainland and other Caribbean island populations (21) (Figure 1B.5). These morphological differences between Cuban and mainland bobwhites cast doubt on the initial hypothesis of recent human-mediated introduction given how little time had elapsed for phenotypic differences to accrue (21, 22). However, German-Cuban naturalist Juan Gundlach published a conflicting account describing the introduction of bobwhites near Havana by a Spanish colonel during the late 18th century, although Gundlach did not know the source of the introduced population. Gundlach was also confused by the degree of morphological differentiation he observed between Cuban and mainland populations of bobwhites (22). Gundlach was able to integrate these competing ideas of recent introduction yet significant differentiation with his theory that Cuban bobwhites were native to the savannas of Pinar del Río province in western Cuba while the translocation of these native individuals by the Spanish colonel around Havana explained the eastward expansion of Cuban bobwhites during the latter half of the 19th century.

To further complicate matters, translocations of Florida bobwhites (*C. v. floridanus*) to mainland Cuba began during the late 19th century (14, 23), and translocated individuals began hybridizing successfully with “native” Cuban bobwhites despite differences in their natural habitats (23). As of the early 20th century, few “pure” Cuban bobwhites were thought to remain outside of Pinar del Río province and Isla de la Juventud (Figure 1) where there were no documented introductions; thus, these two localities have been suggested as the last refuges of “pure” Cuban bobwhites (17). However, based on the similarities in plumage between the “pure” Cuban bobwhites from Isla de la Juventud and bobwhites from the Caribbean slope of southern Mexico (including the states of Veracruz and Tabasco; Figure 1), it has also been suggested that Cuban bobwhites were introduced from southern Mexico during Spain’s colonization of both countries (24).

Collectively, this prior work suggests the following hypotheses: (1) bobwhites were introduced to Cuba from the southeastern USA during the Spanish colonial period (20); (2) bobwhites were endemic to western Cuba prior to the arrival of Europeans (22); and (3) bobwhites were introduced to Cuba from southern Mexico during the Spanish colonial period (24). Similar to other studies where natural and human-mediated dispersal events may have interacted over relatively recent timescales, molecular efforts to demystify the origins of Cuban bobwhites have produced equivocal results. For example, an analysis of two mitochondrial markers found shared haplotypes between eight bobwhites collected from central and eastern Cuba and individuals collected from Mexico and the USA. However, these data failed to identify any unique Cuban haplotypes; showed little resolution in the trees and haplotype networks amongst Cuban, Mexican, and USA bobwhite populations; and showed discordance between haplogroups and population boundaries (25). A subsequent analysis using thousands of ultraconserved element (UCE) loci placed Cuban bobwhites sister to a clade of individuals collected from southern Mexico – suggesting the two populations share a common ancestor (26). Although this analysis only sampled a single individual from Cuba, the specimen was collected in Pinar del Río province – one of the potential remaining refuges of “pure” Cuban bobwhites (17, 22). While these results suggest that Cuban bobwhites share genetic ancestry with multiple mainland bobwhite populations, the origin and timing of how bobwhites arrived to and dispersed across Cuba remains unclear.

To address these questions, we used a target capture approach appropriate for historical specimens (RADcap; (27)) to collect thousands of restriction-site associated (RAD; (28–30)) loci from 109 bobwhite specimens sampled during the 1850s to the 1960s throughout mainland Cuba, Isla de la Juventud, and source populations in the USA and Mexico. Using phylogenetic, population genetic, and demographic analyses in concert with historical records, we evaluated the three hypotheses that have been proposed to explain the origin of Cuban bobwhites.

## Results

### Sequence data from tissue and toepad samples

Sequence data collected for both tissue and toepad samples were comparable, although we observed some differences between sample types and greater variance in all metrics among sequence data collected from toepads (Table S2). Specifically, we collected an average of 1,629,289 reads (range 206,636 - 2,483,216, 95% confidence interval (CI) ± 377,407) from tissue samples prepared as 3RAD libraries, and 1,338,505 reads (range 269,012 - 5,980,941, 95% CI ± 158,242) from toepad samples prepared as standard genomic libraries. The main difference between the two sample types was the efficacy of target capture: 95.7% of reads collected from tissue samples were on-target, whereas 24.1% of reads were on-target for toepad samples. We also discarded substantially more duplicate reads from toepad samples (24.8%) than from tissues (4.96%). After removing duplicates, the average depth of coverage across targeted SNPs was lower in tissues (range 35-72, mean 47 ± 4 95% CI), although the variance was greater for toepads (range 19-209, mean 66 ± 6 95% CI).

After filtering (*Materials and Methods*), the Set1 VCF files used for population genetic and phylogenetic analyses contained 1,258 SNPs (Set1a, ingroup only) and 1,267 SNPs (Set1b, ingroup+outgroup), and the Set2 VCF file used for demographic modeling contained 2,228 SNPs.

### Population structure & genetic diversity

sNMF runs for individuals from the USA, Mexico, and Cuba identified four populations as the bestfitting value of K, a result that is broadly concordant with sampling region. All Cuban and USA samples were assigned to their own populations, respectively, and the Mexican samples were assigned to either a northern or a southern population, consistent with phylogenetic breaks observed in UCE data for this species that correspond to the Transvolcanic Belt (26, 31, 32) (Figure 2A). sNMF also showed that the four populations were well-differentiated with little evidence of admixture in most individuals. The exceptions to this general trend were admixture between the northern and southern Mexico populations evident in samples collected from the Caribbean slope of southern Mexico and admixture between the northern Mexico population and USA individuals collected in Louisiana and Georgia. Notably, individuals from the Caribbean slope of southern Mexico shared alleles with Cuban individuals (Figure 2A). The DAPC results also identified four populations as best fitting the data, and population assignments were identical to the results from sNMF (Figure 2B). PCA analyses, which did not require *a priori* selection of the number of clusters in the data, also showed four well-differentiated populations and identical population assignment to sNMF and DAPC results (Figure 2C). When we included Black-throated Bobwhites (*Colinus nigrogularis*) as the outgroup in similar analyses, the best-fitting K-value suggested by sNMF increased to five populations, with little change in the admixture proportions estimated among the bobwhite samples or in the clustering results from either the dAPC or PCA (Figure S1). These results suggest Black-throated Bobwhite populations from the Yucatán Peninsula were not the source of Cuban bobwhites.

**Figure 2.**
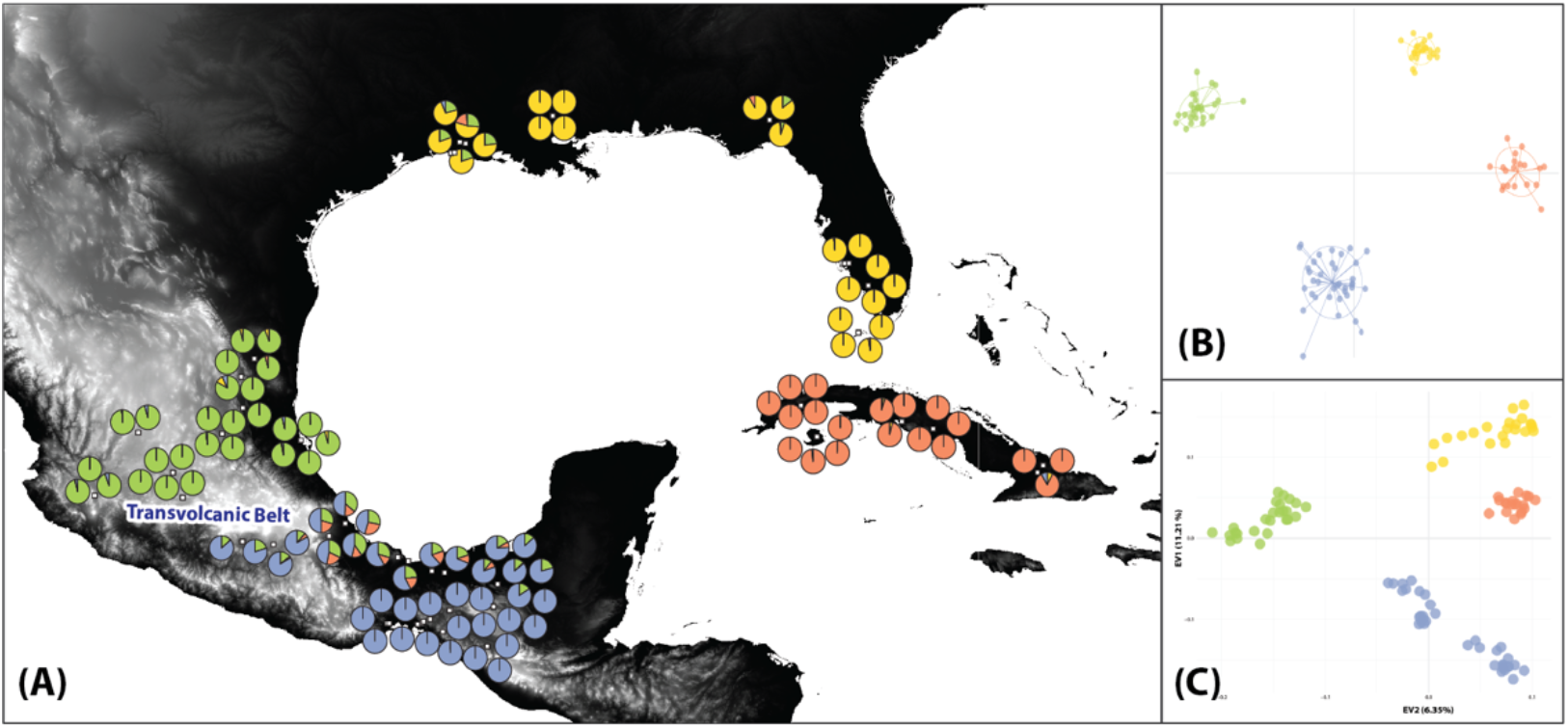
Population structure of Northern Bobwhites from Cuba, the USA, and Mexico. **(A)** sNMF results for the best-fitting K-value of four populations. Pie charts show admixture proportions for each individual plotted by sampling locality (points have been jittered around exact coordinates to allow easier viewing). **(B)** DAPC results for best-fit four population clusters: USA (yellow), northern Mexico (green), southern Mexico (blue), and Cuba (orange). **(C)** PCA results for the same set of individuals. Clusters have been colored as in (A-B) to reflect population assignment from sNMF and DAPC.

Across all populations, mean nucleotide diversity was 0.277 (0.268 - 0.285 95% CI), and observed heterozygosity was lower than expected (t = 21.845, df = 5067, p-value = 2.2e-16) (Table S3). When we considered populations separately, nucleotide diversity and observed and expected heterozygosity were highest among USA bobwhites and lowest among southern Mexico bobwhites (Table S3). All four populations had positive inbreeding coefficients, ranging from 0.050 in the Cuban population to 0.155 in southern Mexico (Table S3). F_ST_ values were moderate between populations (Table S4), with the Cuban population showing the least differentiation compared to the USA, followed by comparisons of Cuban individuals to individuals collected from the northern Mexico and southern Mexico populations (Table S4). We identified private alleles in all four populations, including seven alleles unique to the Cuban population (Table S4). When we expanded the analysis to look for alleles unique to pairs of populations, we found one private allele shared between Cuba and northern Mexico, 11 shared between Cuba and southern Mexico, and 47 shared between Cuba and the USA (Table S4).

### Phylogenetic analyses

SNAPP analyses using a broad geographic subsampling scheme, in which we randomly sampled individuals from each of the four populations identified by sNMF (Cuba, northern Mexico, southern Mexico, and the USA), produced different species tree topologies (Figure 3). In four of the five analyses, the consensus topology resolved the Cuban population as sister to the USA population, with Cuba+USA resolved as sister to a clade consisting of both Mexican populations (Figure 3A-D). However, the remaining analysis resolved the Cuban population as sister to the two Mexican populations (Figure 3E). Across all five analyses, the posterior distribution of trees showed considerable variation in topology, including allele sharing between the Cuban population and multiple mainland populations (Figure 3A-D).

**Figure 3.**
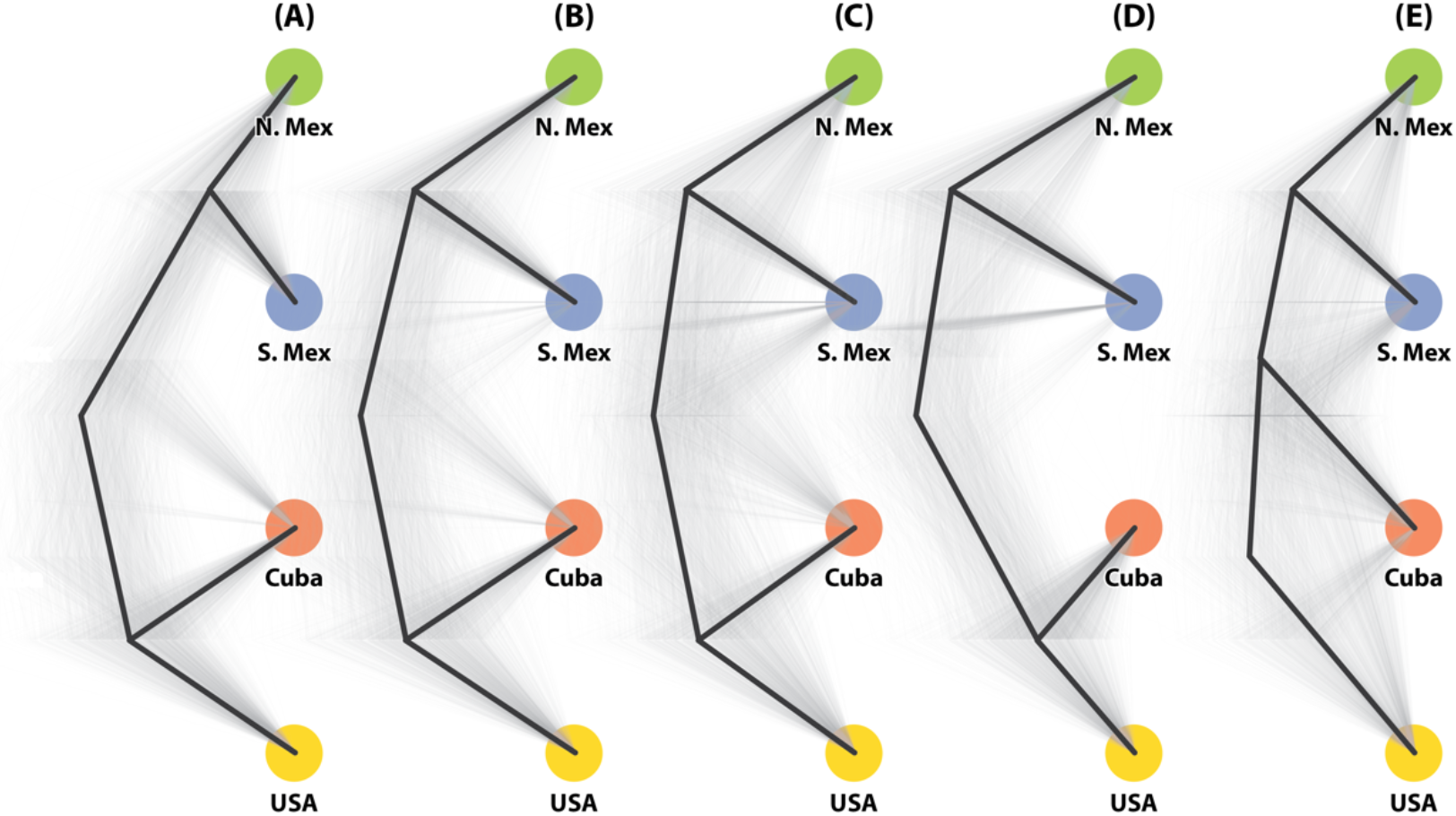
Evolutionary relationships of Northern Bobwhite populations from Cuba, the USA, and Mexico. Species trees from five independent SNAPP analyses using five randomly sampled individuals per population **(A-E).** Consensus topology is depicted by black lines; alternate topologies are depicted using gray lines. The samples used for each analysis are listed in Table S7.

SNAPP analyses with a finer population-level subsampling scheme, which we used to understand the complicated history of introductions across Cuba, suggested that there was a geographic pattern in the discordance we observed in the first subsetting scheme: individuals collected in Pinar del Río, the westernmost population in Cuba, were resolved as sister to the southern Mexico population, with northern Mexico and USA populations forming the sister clade to this group (Figure 4A). However, individuals collected from three populations in the central and eastern provinces of Cuba, as well as individuals collected from Isla de la Juventud, were resolved as sister to the USA population, with northern and southern Mexico populations forming a subsequent sister clade (Figure 4B-E). As in the first subsetting scheme and despite the differences in topology, the posterior distribution of trees in all analyses showed allele sharing between the Cuban population and multiple mainland populations.

**Figure 4.**
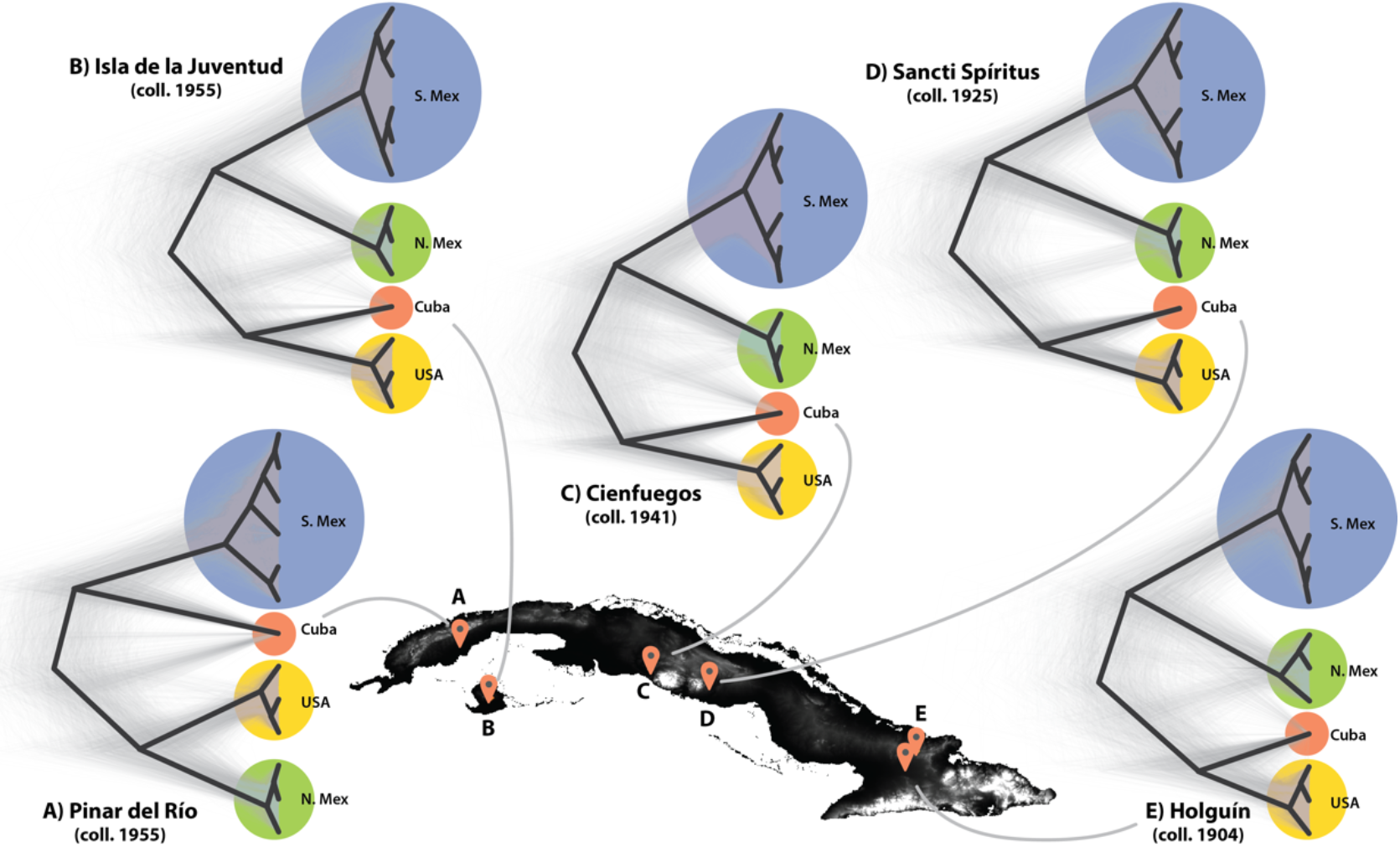
Cuban bobwhite populations have heterogeneous evolutionary histories. Species trees from five independent SNAPP analyses using 3-5 samples per Cuban population and two randomly sampled individuals from all subpopulations (Table S1) of each mainland population **(A-E).** The year the Cuban samples were collected is listed below the locality name. Consensus topology is depicted by black lines; alternate topologies are depicted using gray lines. The samples used for each analysis are listed in Table S8.

### Demographic analyses

To allow objective comparisons of historically and biologically informed alternative evolutionary scenarios for the arrival of bobwhites to Cuba, we tested 45 demographic models (*Materials and Methods).* In the initial comparison of all 45 models, only three models fell within the candidate set. After running each of these three models 100 times to search the likelihood space, only two models remained in the confidence set with Akaike weights (*w_i_*) of 0.503 and 0.493 (Table S5; Table S11).

Both models support a scenario in which the Cuban bobwhite population was founded by bobwhites from southern Mexico, followed by introduction of bobwhites from the USA (Figure 5; Table S5). The model receiving greater support (*w_i_* = 0.503) constrained the initial colonization from southern Mexico between 0.5-5.0 kya (estimated at ~855 years ago or ca. 1165 CE), with the introduction from the USA occurring ~300 years ago (ca. 1720 CE). The second-best model (*w_i_* = 0.436) constrained the timing of colonization to within the past 500 years and estimated slightly younger ages for founding and migration events (colonization from southern Mexico ~467 years ago, introduction from the USA ~158 years ago). Both models estimated small effective population sizes for Cuban bobwhites (393-752 individuals), consistent with a scenario of a small founding population. Both models estimated the pulse probability from the USA at ~58%, meaning 58% of the effective population size of Cuban bobwhites are migrants from the USA (33). Remaining parameter estimates were concordant between the two models (Table S5). Both models estimated: (1) the divergence between Northern Bobwhites and Black-throated Bobwhites occurred ca. 1.563 Ma, which was equal to the constraint we imposed for this event based on results from a time-calibrated phylogeny (34); and (2) simultaneous divergence of the three mainland lineages ca. 334 kya. The effective population size estimates were similar for the three mainland bobwhite lineages, ranging from ~920,000 for the USA to ~1.1 million for southern Mexico and ~1.3 million for northern Mexico. Effective population size of Black-throated Bobwhites was estimated at ~2.2 million (Table S5).

**Figure 5.**
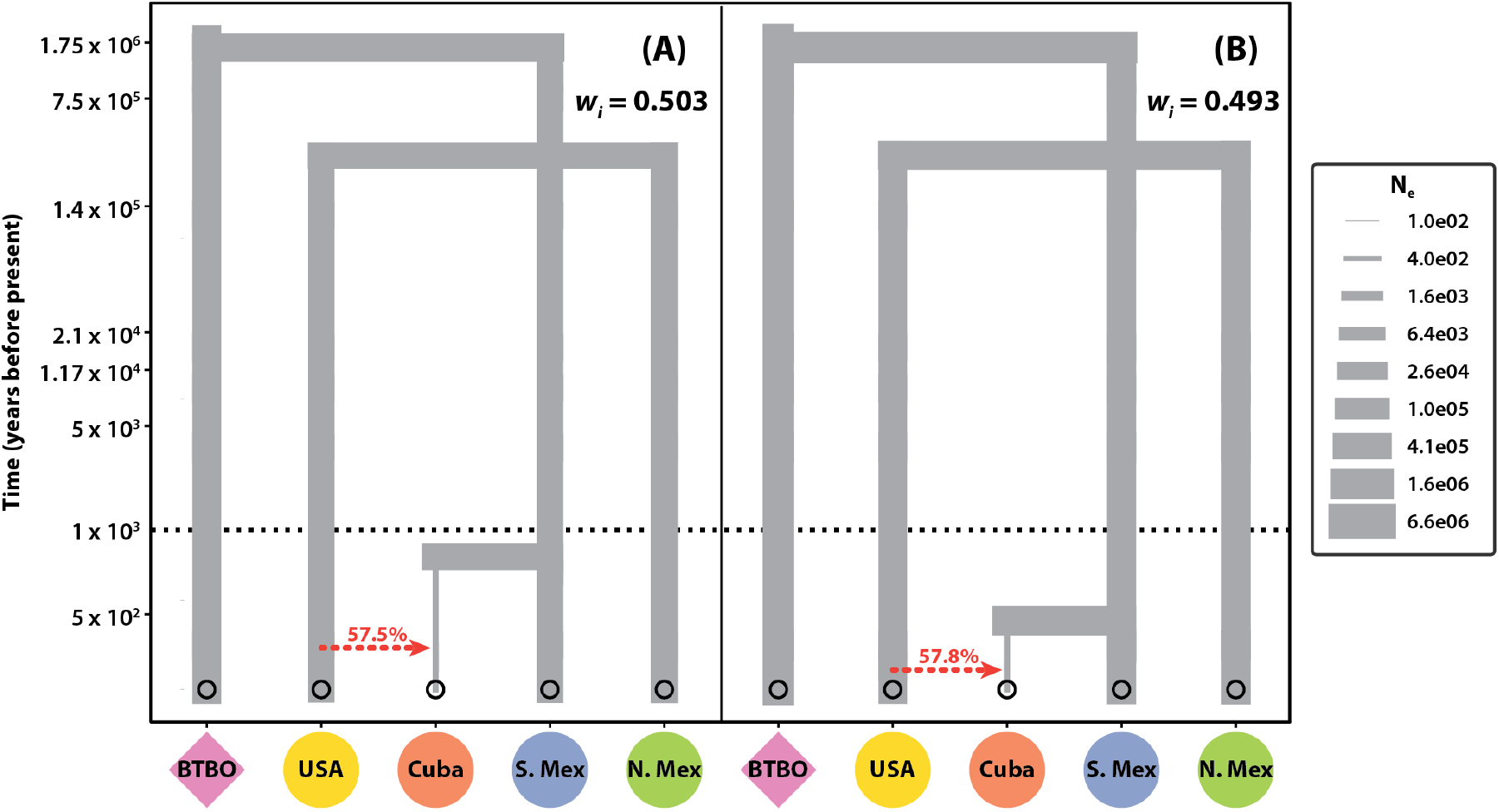
Parameter estimates of the best supported demographic models. **(A-B).** Y-axis is log-scaled above the dotted line. Timing and direction of migration is shown with red-dashed arrows; percentages represent the percent of the Cuban effective population that are migrants from the USA. BTBO = Black-throated Bobwhites; S. Mex = southern Mexico; N. Mex = northern Mexico. Branches are scaled for effective population size (N_e_); scale shown at right.

## Discussion

### Cuban bobwhites are a unique population born through hybridization of divergent lineages

Together, our data suggest that a small number of bobwhites from the Caribbean slope of southern Mexico colonized Cuba, followed by significant introgression from bobwhites introduced to Cuba from the southeastern USA. The conflicting signals of ancestry present in the SNAPP analyses suggest that Cuban bobwhites are the result of interacting evolutionary lineages, consistent with introductions from mainland source populations in southern Mexico and the USA (Figure 3, Figure 4). Our demographic model results are unequivocal that the Cuban bobwhite population was founded by individuals from southern Mexico, followed by substantial introgression with bobwhites introduced from the southeastern USA (Table S5; Figure 5). Models supporting an alternate order of introduction received no support (Table S11).

Although the sNMF results showed little evidence of allele sharing between Cuban bobwhites and individuals in northern Mexico and the USA, the nine individuals that we sampled from the Caribbean slope of southern Mexico showed a considerable degree of allele sharing with the Cuban population (11-26% population assignment) (Figure 2). This finding is consistent with a scenario in which individuals from this region of southern Mexico founded the Cuban population, followed by subsequent founder effects and genetic drift in the Cuban population which can significantly alter allele frequencies without much effect on heterozygosity (35). Despite the small effective population sizes for Cuban bobwhites estimated by the demographic models (Figure 5, Table S5), we observed higher heterozygosity and allelic diversity among Cuban bobwhites than in the much larger southern Mexico population (Table S3), consistent with a founding population bottleneck followed by substantial movement of individuals from the USA into Cuba, similar to patterns of genetic diversity in recently introduced populations of Caribbean anoles (36, 37) and frogs (13). Taken together, these results reject the hypothesis that Cuban bobwhites were originally introduced from the southeastern USA, although our demographic model parameter estimates suggest that the effective size of the Cuban bobwhite population comprises more individuals from the USA than from southern Mexico (Table S5).

Conversely, SNAPP results support conjecture (17, 22) that Pinar del Río province is a reservoir of “native” Cuban bobwhites and potentially the site of their founding population (Figure 4). Pinar del Río was the only population for which SNAPP inferred a species tree that resolved Cuban bobwhites as sister to bobwhites collected in southern Mexico, rather than the USA (Figure 4A), unlike the three populations in central and eastern Cuba where 19th century introductions from the southeastern USA have been documented (23). However, our SNAPP results challenge the idea that Isla de la Juventud remains a population of “native” Cuban bobwhites that have not experienced subsequent introgression with bobwhites from the USA. Individuals from this locality were resolved as sister to the USA population rather than sister to the southern Mexico population (Figure 4B). That said, given the small size of the island (2,200 km^2^) and our small sample size (n=5), a single additional individual could conceivably alter this result.

In effect, the Cuban bobwhites we sampled (collected between 1859-1966) are advanced generation hybrids between two well-differentiated lineages of Northern Bobwhites that have been evolving somewhat independently for >300,000 years (Figure 5, Table S5). This conclusion explains observations of diverse plumage phenotypes in Cuban bobwhites, with specimens collected from around Cuba and Isla de la Juventud displaying plumage typical of Florida bobwhites (38), Caribbean slope bobwhites (24), apparent Cuban-Floridian hybrids (23), and the distinct *Colinus cubanensis* plumage described by Gould (21) (Figure 1B).

### The timing of bobwhite arrival on Cuba suggests human-mediated introduction

Although both demographic models in our confidence set agree on the source and order of introductions to Cuba, the small difference in their Akaike weights suggests they are equivocal regarding the precise timing of these events. The best-weighted model suggests bobwhites arrived in Cuba from southern Mexico 855 years ago (ca. 1165 CE), but this scenario is only 1.02 times more likely given our data than the model that estimated the arrival of bobwhites 467 years ago (ca. 1553 CE). This equivocacy may be the result of how we constrained our models. In both candidate models, the divergence time between Northern Bobwhites and Black-throated Bobwhites was estimated to be slightly younger than the older limit that we allowed based on a previous time-calibrated phylogeny. We constrained this split to the older 95% HPD from that analysis, which is ~400,000 years older than the mean divergence time of 1.133 Ma (34) – suggesting that without this constraint, the divergence time estimates from our dataset could be considerably older than that of previous studies. Our choice of mutation rate and generation time could also have influenced these estimates, so we caution over-interpreting the precise dates obtained from demographic model runs. Ultimately, the equivocacy between the two models in the confidence set constitutes a difference of a few hundred years and resolving this exact difference may go beyond the degree of resolution we can expect from this type of molecular data.

The demographic model results unambiguously indicate that bobwhites first arrived in Cuba very recently, well after the earliest documented human settlement. The older end of our divergence time estimates suggest that bobwhites were already present in Cuba at the time of European settlement beginning in 1510 (Table S5); however, this timing does not preclude human involvement in the introduction of bobwhites to Cuba. The oldest archeological evidence of human settlement on Cuba dates to 3100 BCE, with subsequent waves of migrants arriving from the Lesser Antilles during the following two thousand years (39). The indigenous practice of moving wildlife, particularly animals used for food, has been well documented in the Caribbean (40, 41), including recent molecular and fossil evidence that Lucayan people shaped the distribution of hutias (*Geocapromys ingrahami*) throughout the Bahamian archipelago (42), although the extent of this practice by indigenous peoples inhabiting Cuba is unclear. There is some speculation that the Taíno people of Cuba reached south Florida during the late 15th century prior to European arrival, and that they may have made contact with the Yucatán Peninsula, although archeological evidence for these conclusions is lacking (reviewed in (43)). The older time frame we inferred for the original introduction of bobwhites to Cuba is consistent with the confusing scenario proposed by Gundlach: that bobwhites were already present on the island prior to Spanish colonization and that bobwhites were introduced near Havana by a Spanish colonel during the late 1700s (22).

The younger divergence time estimates from our demographic models place the introduction of bobwhites to Cuba shortly after European colonization. Cuba was one of the islands visited by Christopher Columbus in 1492, and the first Spanish settlers arrived during 1510, establishing permanent settlements in present-day Havana by 1515 (44, 45). Between 1575-1779, the main trans-Atlantic shipping artery used by Spanish fleets traveled between Veracruz and Havana before departing across the Atlantic (Figure 1), with as many as 100 ships making this journey each year by the end of the 16th century (reviewed in (46)). This history is remarkably consistent with our estimates, which indicated the introduction of bobwhites occurred during the mid-16th century (Figure 4, Table S5). The historical Veracruz-Havana connection also supports the signals of shared ancestry in our sNMF analyses between bobwhites in Cuba and the Caribbean slope of southern Mexico (Figure 2) and the previously reported sister relationship between Cuban bobwhites and a clade including Caribbean slope populations (26). These results are consistent with the hypothesis that bobwhites were introduced to Cuba from the southern Caribbean slope of Mexico prior to the 19th century (24). Furthermore, the younger dates we estimated suggest that bobwhites from the southeastern USA arrived in Cuba ca. 1860, which is supported by the collection of Cuban bobwhites with Florida-like plumage during the following decades (23).

Based on our results, we cannot reject either Gundlach’s hypothesis that bobwhites were already present in Cuba prior to European arrival or Parkes’ hypothesis that bobwhites were brought to Cuba from Mexico by the Spanish, in part because they are not mutually exclusive. The range of introduction dates estimated by our demographic analyses could support Gundlach’s theory that there was already a population of bobwhites in Cuba when the Spanish began introducing them, while the younger dates are consistent with the hypothesis that the Spanish were wholly responsible for the introduction of bobwhites. Gundlach did not specify any sources of bobwhites introduced to Cuba, whereas Parkes was unequivocal that Cuban bobwhites were introduced from the Caribbean slope of Mexico. The same shipping vessels traveling between Veracruz and Havana also stopped in Florida (46), where the Spanish established colonies beginning in 1568, so a Floridian origin of Cuban bobwhites introduced by the Spanish is plausible within this timeframe. However, our results clearly support southern Mexico as the original source of the Cuban bobwhite population.

In the absence of explicit documentation that humans (either indigenous or Europeans) brought the first bobwhites to Cuba from southern Mexico, our data cannot exclude a scenario in which bobwhites arrived on Cuba through some kind of natural dispersal; however, following established criteria for distinguishing translocated versus naturally occurring populations of animals (10), we contend that it is extremely unlikely, given both the dispersal capability of bobwhites and the evidence from our genetic data. Bobwhites are largely sedentary and rarely disperse over distances greater than a few kilometers (15, 16). The shortest distances between Cuba and mainland North America are over 200 kilometers to either south Florida (~225 km) or the Yucatán peninsula (~210 km), and the distance between the Caribbean slope and Cuba is even greater (~1,290 km). Furthermore, many sedentary birds are averse to dispersal over water (47), making a natural dispersal scenario improbable. The only other odontophorid quails found on islands are Catalina California Quails (*Callipepla californica catalinensis*), which are considered endemic to Catalina Island off the coast of Southern California; notably, this population is suspected to have been introduced by Paleoindians following their arrival to the island ~12 kya (48, 49). Finally, our estimate of a recent introduction is consistent with the lack of fossil evidence of bobwhites (or any galliformes) on Cuba during the Quaternary, despite a diverse avian fossil record (50, 51) and the identification of several bobwhite fossils from Florida (52). In summary, the population genetic and demographic evidence we report here, combined with historical records and observations, supports the conclusion that bobwhites arrived in Cuba through human-mediated introduction within the recent past.

### Conclusions

In light of our results, it is worth reconsidering Gould’s taxonomic description of Cuban bobwhites (21). Gould’s recognition of Cuban bobwhites as distinct from all other mainland populations of bobwhites (including *C. v. pectoralis* of Veracruz, which he also described), is largely supported by our data. Despite their recent arrival on the island, the combination of genetic drift, multiple introductions, and small population size have given Cuban bobwhites a distinct allelic profile that is well-differentiated from the mainland bobwhite populations we sampled (Figure 4, Tables S3-S4). The unique plumage phenotypes observed in Cuban bobwhites, which inspired Gould’s taxonomic description, also suggest a compelling avenue of future study: as hybrids between two phenotypically divergent lineages, Cuban bobwhites offer a unique opportunity to study the genetic basis of plumage traits that have given rise to the remarkable phenotypic diversity within bobwhites, which have more subspecies (described by male plumage) than 99% of all other birds (53).

The combination of historical specimens and a wealth of historical records provided a unique opportunity to test demographic hypotheses in this system, but our results have implications beyond bobwhites. Many animal distributions have been shaped by human activity over thousands of years, indirectly through landscape and habitat changes as well as through direct introduction and translocation of animals. Yet, in some cases the extent of human involvement or our ability to detect human effects remains unclear. Our results demonstrate that genomic data from historical specimens are sufficiently powerful to infer complex population histories over very recent timescales, and similar approaches could be applied to study other, ambiguous animal distributions, particularly on islands.

## Materials and Methods

### Sampling design and DNA extraction

We sampled 109 individuals, including 95 toepads from historical museum specimens collected between 1859 - 1966 (median collection year = 1941) and 14 tissue samples collected between 1985 - 2014 (median collection year = 2010; Table S1). We included 104 Northern Bobwhites from: Cuba (n=19), the USA (n=22), and Mexico (n=63), as well as five Black-throated Bobwhites (*C. nigrogularis*) from the Yucatán peninsula (Table S1), which we used as outgroup individuals in some analyses (26). We extracted total DNA from tissues using a Qiagen DNeasy Blood & Tissue Kit following the manufacturer’s instructions, and we extracted total DNA from toepads using a phenol-chloroform protocol (54). Following extraction, we quantified samples with a Qubit Fluorometer (Life Technologies, Inc.).

### Library preparation, targeted enrichment, and sequencing

For tissue samples, we prepared three 3RAD pools containing eight uniquely indexed individual libraries that were constructed in a way that allowed us to identify and remove PCR duplicates. For toepad samples, we prepared standard genomic libraries using the KAPA Hyper Prep library preparation kit (F. Hoffmann-La Roche AG) and custom-indexes (55). Prior to enrichment, we quantified libraries using a Qubit Fluorometer and combined toepad libraries into pools of eight individuals at equimolar ratios. We enriched all library pools using our custom RADcap bait set following the myBaits Hybridization Capture for Targeted NGS manual v4.01 and sequenced all pools using two partial lanes of 150-bp paired-end (PE150) sequencing on an Illumina NovaSeq 6000 (Novogene, Sacramento, CA). Remaining samples on each lane had non-overlapping indexes. Additional details regarding bait design, library preparation, targeted enrichment, and sequencing are provided in the Supplemental Methods.

### SNP calling & filtering

Because tissue and toepad libraries were prepared with different types of indexes, we performed slightly different steps to demutiplex sequencing reads, trim reads for adapters and low-quality bases, and remove PCR duplicates. The full procedure is described in the Supplemental Methods. We aligned reads to the bobwhite reference genome (56) using bwa v0.7.17 (57) and samtools v1.10 (58), and we called SNPs using a parallel implementation of the Best Practices for Variant Discovery (59, 60) in GATK v4.1.9.0, including base quality score recalibration (61).

Because the analyses we implemented assume sites are putatively neutral and unlinked (33, 62), we used VCFTools to filter SNPs in the recalibrated VCF file to the single locus targeted during bait design. We also used VCFTools to exclude: loci determined to be indels, sites with less than a minimum depth of 10, sites with more than 10% missing data, and sites out of Hardy-Weinberg equilibrium due to excess heterozygosity, which can indicate genotyping errors (63). Because missing data can bias population structure assignment (64), we also removed individuals missing more than 25% of calls at the remaining sites.

Because particular filters applied to a set of variable SNPs can affect different types of analyses in unexpected ways (65), we produced two types of VCF files: Set1 for population genetic summary statistics, genetic structure, and phylogenetic analyses; and Set2 for demographic analysis. To produce the first set of files (Set1), we used VCFTools to remove loci having a minor allele count (MAC) less than three (66), and we output files: (Set1a) including all bobwhite samples + the outgroup individuals (ingroup+outgroup); and (Set1b) including only the Cuban, USA, and Mexican populations of bobwhites (ingroup only). The inclusion of the outgroup individuals in the Set1a data file allowed us to test where there was any allele sharing between Black-throated Bobwhites and Cuban bobwhites because some populations of Black-throated Bobwhites on the Yucatán Peninsula are physically closer to Cuba than populations of Northern Bobwhites in either Florida or the Caribbean slope of southern Mexico. To produce the second file for demographic analyses (Set2), we removed the MAC filtering because rare alleles can be important for inferring demographic history (67), and we included all bobwhites + the outgroup individuals so that we could include an external constraint on the common ancestor of the ingroup and outgroup (see below). Additional details on base quality score recalibration, SNP calling, and filtering procedures are provided in the Supplemental Methods.

### Patterns of population genetic diversity

To assess population structure and potential admixture among populations, we calculated ancestry coefficients in sNMF (62) using Set1a and Set1b VCF files. To further characterize the patterns of genetic clustering in our dataset, we performed a discriminant analysis of principal components (DAPC) using VCFR v1.12.0 (68) and adegenet v2.1.3 (69) in R v4.0.3 (70). Finally, we performed a principal components analysis (PCA) using SNPRelate (71) in R v4.0.3 (70) because PCA does not require *a priori* selection of the number of clusters present in the data.

To describe patterns of differentiation and genetic diversity within and among bobwhites (ingroup only), we calculated population genetic summary statistics (observed and expected heterozygosity, allelic richness, inbreeding coefficients, nucleotide diversity, and F_ST_ between each pair of populations) using the Set1b VCF file. We also calculated the number of private alleles in each population using a custom Python (72) script, which can be found on GitHub (73). Additional details on analyses of genetic structure and population genetic summary statistics are in the Supplemental Methods.

### Phylogenetic analyses

To investigate the evolutionary relationships among populations, we used the Set1b VCF file to estimate species trees using SNAPP v1.5.1 (74) implemented in Beast v2.6.3 (75). We chose SNAPP because it was designed to model the coalescent process using allele frequencies from biallelic SNP data and because it produces output that enables users to visualize conflicting signals due to allele sharing among multiple populations which we expected to exist in our dataset based on the possibility of multiple introductions into Cuba. Because SNAPP is computationally intensive, precluding us from estimating a species tree for our entire dataset, we used two types of subsampling schemes to provide resolution at both broad and fine geographic scales: 1) five replicate subsamples containing five randomly sampled individuals from each of the four geographic populations identified by our sNMF analyses (USA, northern Mexico, southern Mexico, and Cuba) (20 individuals per analysis) (Table S7); and 2) locality-specific subsets for each of the five, distinct Cuban localities that included all individuals from that location (range 3-6 individuals) as well as two randomly selected individuals from each of the twelve mainland populations representing northern Mexico, southern Mexico, and USA (total 27-30 individuals per analysis) (Table S7).

We ran each SNAPP analysis with default parameters for 2 million iterations, sampling every 1,000 iterations and discarding the first 10% of sampled iterations as burn-in. We used Tracer v1.7.2 (76) to visualize estimated parameters and to ensure that the effective sample size of each (ESS) was greater than 200 (77). We visualized the resulting species trees and posterior distribution of gene trees using DensiTree v2.2.7 (78, 79). Additional details on random sampling protocols and SNAPP analyses are provided in the Supplemental Methods.

### Demographic analyses

Inferring evolutionary relationships and visualizing allele sharing among populations provide one mechanism for understanding population origins, but these types of analyses do not typically allow objective comparisons of alternative evolutionary scenarios, including comparisons of the timing of evolutionary events and/or the source(s) and direction of migrants to specific populations. This is particularly true in the absence of suitable fossil calibrations, which is currently the case for Cuban bobwhites (50). To perform these types of analyses, we used momi2 (33) with the site frequency spectrum (SFS) derived from the Set2 VCF file.

Because the historical record contradicts a hypothesis of a single introduction of bobwhites to Cuba (17, 23), we tested two categories of models given our data: single source models with a second dispersal event and multiple source models. We designed the single source models to test scenarios where one of the mainland populations (the ancestor of northern + southern Mexico, northern Mexico, southern Mexico, or USA) founded the Cuban population, followed by a single dispersal event from the same founding population to Cuba. This design is consistent with either natural or human-mediated dispersal scenarios from a single geographic origin. We designed the multiple source models to test scenarios where one of the mainland populations founded the Cuban population, followed by a single dispersal event from a different mainland population during the last 500 years. This design is consistent with either natural or human-mediated dispersal scenarios establishing the Cuban population followed by human-mediated introductions during the last 500 years, as noted in several historical sources (17, 22, 23). We constrained models in the multiple source category to always include the USA, either as the founding population or as the source of more recent dispersal to Cuba, because SNAPP analyses always showed allele sharing between bobwhite populations in the USA and Cuba. For example, we did not model a scenario in which northern Mexico bobwhites founded the Cuban population, followed by recent dispersal of bobwhites from southern Mexico because this was inconsistent with the SNAPP results. To allow tests of hypotheses regarding the timing of bobwhite arrival on Cuba, we incorporated five different temporal scenarios to all models in both categories for the founding of the Cuban population – specifically, that the Cuban bobwhite population was founded: 1) within 0.5 kya (since European arrival on Cuba; (44); 2) between 0.5-5.0 kya (since Indigenous arrival on Cuba; (39)); 3) between 5.0-11.7 kya (since the Pleistocene-Holocene Transition; (80)); 4) between 11.7-23.0 kya (since the last glacial maximum; (80)); or 5) between 23.0-140.0 kya (since the penultimate glacial maximum; (81). In total, this produced 45 separate models (Figure S2; Table S9).

Before running momi2, we set the effective size of the population ancestral to bobwhites and their sister species, Black-throated Bobwhites to 3.35 x 10^5^, the mean value estimated across multiple G-PhoCs runs (additional details on G-PhoCs analyses are in the Supplemental Methods); we specified a generation time of 1.22 years, which was the median value estimated from multiple radiotelemetry and survivorship studies of wild bobwhites (82); and we specified a mutation rate of 1.91 x 10^-9^ sites per year, which was estimated from chickens (83). Finally, we constrained the divergence time between Northern Bobwhites and Black-throated Bobwhites to have occurred since 1.563 Ma, which was the older bound of the 95% highest posterior density interval estimated for their divergence in a time-calibrated phylogeny (34).

We ran each of the 45 models 10 times and computed corrected Akaike information criterion (AICc) scores (84, 85) for all models in each set of the 10 model runs. We then used AIC-based model comparison (86) to rank and compare models (Table S10). For each of the three models that fell within the confidence set (87) among any of the 10 runs, we completed an additional 90 runs. We selected the best (highest) log-likelihood value for each model out of 100 (10+90) runs, and we used the best log-likelihood obtained for each of the three top models to compute final AICc scores, delta AICc values, and Akaike weights following Burnham and Anderson (86).

## Acknowledgments

We thank the following museums and collections staff that provided loans for this project: Delaware Museum of Natural History (DMNH - Jean Woods); University of Kansas Biodiversity Research Institute (KU - Mark Robbins); Natural History Museum of Los Angeles County (LACM - Kimball Garrett); Louisiana State University Museum of Natural Science (LSU - Donna Dittmann, Van Remsen, & Fred Sheldon); Harvard Museum of Comparative Zoology (MCZ - Jeremiah Trimble); Moore Laboratory of Zoology (James Maley & John McCormack); Royal Ontario Museum (ROM - Santiago Claramunt); San Diego Museum of Natural History (SDNHM - Phil Unitt); University of Michigan Museum of Zoology (UMMZ - Janet Hinshaw); National Museum of Natural History (USNM - Christopher Milensky); University of Washington Burke Museum (UWBM - Sharon Birks); Western Foundation of Vertebrate Zoology (WFVZ - René Corado); and the Yale Peabody Museum (YPM - Kristof Zyskowski). We would additionally like to thank Christopher Milensky and the National Museum of Natural History for providing access to specimens for photography. We also thank Pete Hosner, who provided the calibration dates for the demographic analyses, and John Carroll, who directed us to Ken Parkes’ work on Cuban bobwhites. Discussions with Anna Hiller, Glaucia Del Rio, and Jessica Oswald provided helpful feedback through the development of this project.

## Data Deposition

Raw sequencing reads will be available from the National Center for Biotechnology Information (NCBI) Sequence Read Archive as part of BioProject PRJNA875956 upon publication. Custom computer code, DNA alignments, analysis inputs, and analysis outputs will be available from the Dryad Digital Repository upon publication.

## Author Contributions

J.F.S., R.T.B., and B.C.F. designed the study. J.F.S. collected and analyzed the data with input from R.T.B. and B.C.F. J.F.S., R.T.B., and B.C.F. wrote the paper. J.F.S., R.T.B., and B.C.F. contributed funds.

## References

1. N. L. Boivin, et al., Ecological consequences of human niche construction: Examining longterm anthropogenic shaping of global species distributions. Proc. Natl. Acad. Sci. U. S. A. 113, 6388–6396 (2016).

2. G. Larson, J. Burger, A population genetics view of animal domestication. Trends Genet. 29, 197–205 (2013).

3. N. D. Tsutsui, A. V. Suarez, D. A. Holway, T. J. Case, Reduced genetic variation and the success of an invasive species. Proc. Natl. Acad. Sci. U. S. A. 97, 5948–5953 (2000).

4. B. A. Mathys, J. L. Lockwood, Rapid evolution of great kiskadees on Bermuda: an assessment of the ability of the island rule to predict the direction of contemporary evolution in exotic vertebrates. J. Biogeogr. 36, 2204–2211 (2009).

5. H. S. Young, I. M. Parker, G. S. Gilbert, A. Sofia Guerra, C. L. Nunn, Introduced Species, Disease Ecology, and Biodiversity–Disease Relationships. Trends Ecol. Evol. 32, 41–54 (2017).

6. C. C. Austin, Lizards took express train to Polynesia. Nature 397, 113–114 (1999).

7. C. Roullier, L. Benoit, D. B. McKey, V. Lebot, Historical collections reveal patterns of diffusion of sweet potato in Oceania obscured by modern plant movements and recombination. Proc. Natl. Acad. Sci. U. S. A. 110, 2205–2210 (2013).

8. C. A. Hofman, T. C. Rick, Ancient biological invasions and island ecosystems: Tracking translocations of wild plants and animals. J. Archaeol. Res. 26, 65–115 (2018).

9. G. R. Summerhayes, Island Melanesian Pasts: A View from Archeology. Genes, Language, & Culture History in the Southwest Pacific (2007).

10. T. E. Heinsohn, Marsupials as introduced species: Long-term anthropogenic expansion of the marsupial frontier and its implications for zoogeographic interpretation. Altered ecologies: Fire, climate and human influence on terrestrial landscapes, 133–176 (2010).

11. T. Heinsohn, Animal translocation:long-term human influences on the vertebrate zoogeography of Australasia (natural dispersal versus ethnophoresy). Aust. Zool. 32, 351–376 (2003).

12. I. Zeisset, T. J. Beebee, Determination of biogeographical range: an application of molecular phylogeography to the European pool frog Rana lessonae. Proc. Biol. Sci. 268, 933–938 (2001).

13. M. P. Heinicke, L. M. Diaz, S. B. Hedges, Origin of invasive Florida frogs traced to Cuba. Biol. Lett. 7, 407–410 (2011).

14. C. B. Cory, A list of the birds of the West Indies, including the Bahama Islands and the Greater and Lesser Antilles, excepting the Islands of Tobago and Trinidad (Estes & Lauriat, 1885).

15. H. L. Stoddard, The bobwhite quail (C. Scribner’s Sons, 1931).

16. G. F. Smith, F. E. Kellogg, W. R. Davidson, W. Mack Martin, A 10-year Study of Bobwhite Quail Movement Patterns. National Quail Symposium Proceedings 2, 6 (1982).

17. T. Barbour, The Birds of Cuba (Memoirs of the Nuttall Ornithological Club, 1923).

18. J. Bond, Origin of the Bird Fauna of the West Indies. Wilson Bull. 60, 207–229 (1948).

19. P. A. Johnsgard, Quails, Partridges, and Francolins of the World (Oxford University Press, 1988).

20. A. D’Orbigny, “Ornothologie” in Historia Fisica, Politica Y Natural de La Isla de Cuba, R. de la Sagra, Ed. (Artus Bertrand, 1839), pp. III–336.

21. J. Gould, A monograph of the Odontophorinæ, or, Partridges of America (John Gould, F. R. S., 1850).

22. J. C. Gundlach, Contribucion á la ornitologia cubana, (Imp. “La Antilla” de N. Cacho-Negrete, 1876).

23. F. M. Chapman, Notes on Birds and Mammals Observed Near Trinidad, Cuba: With Remarks on the Origin of West Indian Bird-life (order of the Trustees, American Museum of Natural History, 1892).

24. K. C. Parkes, The origin of the Cuban Bobwhite “Colinus virginianus cubanensis” (Gray) (1990).

25. D. Williford, R. W. Deyoung, R. L. Honeycutt, L. A. Brennan, F. Hernández, Phylogeography of the bobwhite (Colinus) quails. Wildlife Monographs 193, 1–49 (2016).

26. J. F. Salter, et al., Historical specimens and the limits of subspecies phylogenomics in the New World quails (Odontophoridae). Mol. Phylogenet. Evol. 175, 107559 (2022).

27. S. L. Hoffberg, et al., RADcap: sequence capture of dual-digest RADseq libraries with identifiable duplicates and reduced missing data. Mol. Ecol. Resour. 16, 1264–1278 (2016).

28. M. R. Miller, J. P. Dunham, A. Amores, W. A. Cresko, E. A. Johnson, Rapid and costeffective polymorphism identification and genotyping using restriction site associated DNA (RAD) markers. Genome Res. 17, 240–248 (2007).

29. N. A. Baird, et al., Rapid SNP discovery and genetic mapping using sequenced RAD markers. PLoS One 3, e3376 (2008).

30. J. W. Davey, et al., Genome-wide genetic marker discovery and genotyping using next-generation sequencing. Nat. Rev. Genet. 12, 499–510 (2011).

31. C. J. Marshall, J. K. Liebherr, Cladistic biogeography of the Mexican transition zone. J. Biogeogr. 27, 203–216 (2000).

32. J. J. Morrone, Fundamental biogeographic patterns across the Mexican Transition Zone: an evolutionary approach. Ecography 33, 355–361 (2010).

33. J. Kamm, J. Terhorst, R. Durbin, Y. S. Song, Efficiently inferring the demographic history of many populations with allele count data. J. Am. Stat. Assoc. 115, 1472–1487 (2020).

34. P. A. Hosner, E. L. Braun, R. T. Kimball, Land connectivity changes and global cooling shaped the colonization history and diversification of New World quail (Aves: Galliformes: Odontophoridae). J. Biogeogr. 42, 1883–1895 (2015).

35. F. W. Allendorf, Genetic drift and the loss of alleles versus heterozygosity. Zoo Biol. 5, 181–190 (1986).

36. J. J. Kolbe, et al., Genetic variation increases during biological invasion by a Cuban lizard. Nature 431, 177–181 (2004).

37. J. J. Kolbe, et al., Multiple sources, admixture, and genetic variation in introduced anolis lizard populations. Conserv. Biol. 21, 1612–1625 (2007).

38. Ridgway Robert, Colinus virginianus cubanensis Not a Florida Bird. Auk 11, 324–325 (1894).

39. L. Allaire, “Archaeology of the Caribbean Region” in The Cambridge History of the Native Peoples of the Americas, F. Salomon, S. B. Schwartz, Eds. (Cambridge University Press, 1999).

40. E. S. Wing, “Native American use of animals in the Caribbean” in Biogeography of the West Indies, (CRC Press, 2001), pp. 481–518.

41. L. A. Newsom, E. S. Wing, On Land and Sea: Native American Uses of Biological Resources in the West Indies (University of Alabama Press, 2004).

42. J. A. Oswald, et al., Ancient DNA and high-resolution chronometry reveal a long-term human role in the historical diversity and biogeography of the Bahamian hutia. Sci. Rep. 10, 1373 (2020).

43. C. S. Chard, Pre-Columbian trade between north and south America. Pap. Kroeber Anthropol. Soc. 1, 1–27 (1950).

44. B. de las Casas, Historia de las Indias (Imprenta y litografia de I. Paz, 1877).

45. R. F. Monzote, From Rainforest to Cane Field in Cuba: An Environmental History since 1492 (Univ of North Carolina Press, 2009).

46. A. Lugo-Fernández, D. A. Ball, M. Gravois, C. Horrell, J. B. Irion, Analysis of the Gulf of Mexico’s Veracruz-Havana Route of La Flota de la Nueva España. J. Marit. Archaeol. 2, 24–47 (2007).

47. R. P. Moore, W. D. Robinson, I. J. Lovette, T. R. Robinson, Experimental evidence for extreme dispersal limitation in tropical forest birds. Ecol. Lett. 11, 960–968 (2008).

48. N. K. Johnson, Origin and Differentiation of the Avifauna of the Channel Islands, California. Condor 74, 295–315 (1972).

49. R. M. Zink, D. F. Lott, D. W. Anderson, Genetic Variation, Population Structure, and Evolution of California Quail. Condor 89, 395–405 (1987).

50. J. Orihuela, An annotated list of Late Quaternary extinct birds of Cuba. Ornitol. Neotrop. 30, 57–67 (2019).

51. N. V. Zelenkov, E. S. Belichenko, Dynamics of the Late Quaternary Avifauna of Western Cuba (Based on Material from El Abrón Cave). Dokl. Biol. Sci. 503, 54–57 (2022).

52. J. A. Holman, Osteology of living and fossil New World quails (Aves, Galliformes) (University of Florida, 1961).

53. E. C. Dickinson, J. V. Remsen Jr., Eds., The Howard & Moore Complete Checklist of the Birds of the World, 4th Ed. (Aves Press, Eastbourne, U.K., 2013).

54. W. L. E. Tsai, J. F. Salter, “Phenol Chloroform Extraction (for toepads)” (Moore Laboratory of Zoology, Occidental College; Louisiana State University, 2018).

55. T. C. Glenn, et al., Adapterama I: Universal stubs and primers for thousands of dual-indexed Illumina libraries (iTru & iNext). bioRxiv (2019) https://doi.org/10.1101/049114.

56. J. F. Salter, et al., A Highly Contiguous Reference Genome for Northern Bobwhite (Colinus virginianus). G3 9, 3929–3932 (2019).

57. H. Li, R. Durbin, Fast and accurate short read alignment with Burrows-Wheeler transform. Bioinformatics 25, 1754–1760 (2009).

58. H. Li, et al., The Sequence Alignment/Map format and SAMtools. Bioinformatics 25, 2078–2079 (2009).

59. G. A. Van der Auwera, B. D. O’Connor, Genomics in the Cloud: Using Docker, GATK, and WDL in Terra (2020).

60. J. F. Salter, B. C. Faircloth, “Running GATK in Parallel” (2021) (June 8, 2021).

61. A. McKenna, et al., The Genome Analysis Toolkit: a MapReduce framework for analyzing next-generation DNA sequencing data. Genome Res. 20, 1297–1303 (2010).

62. E. Frichot, F. Mathieu, T. Trouillon, G. Bouchard, O. François, Fast and efficient estimation of individual ancestry coefficients. Genetics 196, 973–983 (2014).

63. B. Chen, J. W. Cole, C. Grond-Ginsbach, Departure from Hardy Weinberg Equilibrium and Genotyping Error. Front. Genet. 8, 167 (2017).

64. X. Yi, E. K. Latch, Nonrandom missing data can bias PCA inference of population genetic structure. Mol. Ecol. Resour. (2021) https://doi.org/10.1111/1755-0998.13498.

65. B. Chattopadhyay, K. M. Garg, U. Ramakrishnan, Effect of diversity and missing data on genetic assignment with RAD-Seq markers. BMC Res. Notes 7, 841 (2014).

66. E. Linck, C. J. Battey, Minor allele frequency thresholds strongly affect population structure inference with genomic data sets. Mol. Ecol. Resour. 19, 639–647 (2019).

67. G. T. Marth, E. Czabarka, J. Murvai, S. T. Sherry, The allele frequency spectrum in genome-wide human variation data reveals signals of differential demographic history in three large world populations. Genetics 166, 351–372 (2004).

68. B. J. Knaus, N. J. Grünwald, vcfr: a package to manipulate and visualize variant call format data in R. Mol. Ecol. Resour. 17, 44–53 (2017).

69. T. Jombart, adegenet: a R package for the multivariate analysis of genetic markers. Bioinformatics 24, 1403–1405 (2008).

70. R Core Team, R: A Language and Environment for Statistical Computing (2020).

71. X. Zheng, et al., A high-performance computing toolset for relatedness and principal component analysis of SNP data. Bioinformatics 28, 3326–3328 (2012).

72. G. Van Rossum, F. L. Drake, Python 3 Reference Manual: (Python Documentation Manual Part 2) (CreateSpace Independent Publishing Platform, 2009).

73. B. C. Faircloth, private-alleles: Compute private alleles in one population relative to another (Github, 2021) (September 7, 2021).

74. D. Bryant, R. Bouckaert, J. Felsenstein, N. A. Rosenberg, A. RoyChoudhury, Inferring species trees directly from biallelic genetic markers: bypassing gene trees in a full coalescent analysis. Mol. Biol. Evol. 29, 1917–1932 (2012).

75. R. Bouckaert, et al., BEAST 2: a software platform for Bayesian evolutionary analysis. PLoS Comput. Biol. 10, e1003537 (2014).

76. A. Rambaut, A. J. Drummond, D. Xie, G. Baele, M. A. Suchard, Posterior Summarization in Bayesian Phylogenetics Using Tracer 1.7. Syst. Biol. 67, 901–904 (2018).

77. A. J. Drummond, S. Y. W. Ho, M. J. Phillips, A. Rambaut, Relaxed phylogenetics and dating with confidence. PLoS Biol. 4, e88 (2006).

78. R. R. Bouckaert, DensiTree: making sense of sets of phylogenetic trees. Bioinformatics 26, 1372–1373 (2010).

79. R. R. Bouckaert, J. Heled, DensiTree 2: Seeing Trees Through the Forest. bioRxiv, 012401 (2014).

80. K. M. Cohen, S. C. Finney, P. L. Gibbard, J.-X. Fan, The ICS international chronostratigraphic chart. Episodes 36, 199–204 (2013).

81. F. Colleoni, C. Wekerle, J.-O. Näslund, J. Brandefelt, S. Masina, Constraint on the penultimate glacial maximum Northern Hemisphere ice topography (≈140 kyrs BP). Quat. Sci. Rev. 137, 97–112 (2016).

82. Y. A. Halley, et al., A draft de novo genome assembly for the northern bobwhite (Colinus virginianus) reveals evidence for a rapid decline in effective population size beginning in the Late Pleistocene. PLoS One 9, e90240 (2014).

83. K. Nam, et al., Molecular evolution of genes in avian genomes. Genome Biol. 11, R68 (2010).

84. C. M. Hurvich, C.-L. Tsai, Regression and time series model selection in small samples. Biometrika 76, 297–307 (1989).

85. C. M. Hurvich, C.-L. Tsai, A corrected Akaike information criterion for vector autoregressive model selection. J. Time Ser. Anal. 14, 271–279 (1993).

86. K. P. Burnham, D. R. Anderson, A practical information-theoretic approach. Model selection and multimodel inference 2 (2002).

87. R. Royall, Statistical Evidence: A Likelihood Paradigm (CRC Press, 1997).

