## Supplemental Material for "An island “endemic” born out of hybridization between introduced lineages"

**This PDF file includes:**

Extended Materials and Methods
Figures S1 to S2
Tables S1 to S11
SI References

### Extended Materials and Methods

#### RADcap study design

Given the shallow relationships among bobwhite subspecies (1–3) and our use of historical museum specimens that ranged in age from 60-170 years, we designed our experiment to collect thousands of SNP loci from each individual using RADcap (4). The RADcap method was designed to collect highly variable RAD loci with the consistency of capture-based sequencing approaches. A second benefit of RADcap is that it enables the collection of RAD data from historical specimens, which often have low quality DNA extracts due to DNA degradation (5) that make standard RAD sequencing difficult (6). In brief, the RADcap process involved collecting an initial dataset using traditional RAD-seq from a limited number of high-quality samples, using these data to design a custom set of baits targeting variable SNPs, and then performing sequence capture with these baits to enrich the targeted set of RAD loci across all samples, both historical and modern (4).

#### 3RAD SNP discovery and bait design

Because we wanted to use our custom bait set for multiple projects within the genus *Colinus*, we collected our initial RADseq dataset from 46 tissue samples encompassing ten subspecies from all four species of *Colinus* (Table S6) following the procedure described in the original RADcap paper (4). We prepared samples using 3RAD (7, 8), a modified version of RADseq (9) that uses two restriction enzymes to cut the DNA and a third restriction enzyme to cut adapter dimers of the phosphorylated adapter, thus improving the efficiency of enzyme digestion and adapter ligation (7). We used *EcoRI-HF* and *XbaI* as the DNA-cutting enzymes, and *NheI-HF* as the adapter-dimer cutting enzyme. We added the enzymes and a unique combination of forward and reverse adapters with short internal indexes to each DNA sample and performed a four-hour digestion at 37° C. Following digestion, we added T4 DNA ligase to the digested DNA and performed adapter ligation. Because each sample was identified by an unique set of indexes, we pooled all 46 samples together and cleaned the post-ligation product with SpeedBeads (10, 11). We added a single combination of iTru5 and iTru7 primers (11) to the pool and performed 12 cycles of PCR (95° C for 2 minutes; 12 cycles of 98° C for 20 seconds, 61° C for 15 seconds, 72° C for 30 seconds; 72° C for 5 minutes), followed by a final SpeedBead cleanup to remove unincorporated primers. We performed size-selection on the pool using a BluePippin (Sage Science, Inc.) centered around 550 bp (±10%), followed by 6 cycles of PCR using Illumina Primer Mix to increase the yield of DNA in our target size range. We used a GeneRead Size Selection Kit (Qiagen, GmbH) to remove unincorporated primers, quantified final pools with a KAPA qPCR quantification kit, and combined pools in equimolar ratios prior to 150-bp paired-end (PE150) sequencing on an Illumina HiSeq 3000 (Oklahoma Medical Research Foundation, Oklahoma City, OK).

The protocol we followed for processing 3RAD data is available on the Faircloth lab website (12). In brief, we used the program *process_radtags* in Stacks v1.48 (13, 14) to demultiplex the sequence data. We aligned the reads to a reference genome assembly of Northern Bobwhite (Cv_LA_1.0) (15) using bwa v0.7.17 (16), and we used samtools v1.8 (17, 18) to filter out unmapped reads, imperfect matches, and reads containing more than five SNPs. We used the Stacks programs *pstacks*, *cstacks*, and *sstacks* to assemble aligned reads into loci, compile a catalog of all loci, and match loci from each sample back to the catalog. Finally, we used the program *populations* to output a variant call format (VCF) file of 171,460 SNPs. We used VCFtools v0.1.13 (19) to filter the dataset so that no more than 5% of allele calls were missing for any loci and for a minor allele frequency (MAF) of >= 0.05. This produced a VCF file of 29,763 SNPs.

Because we wanted our baits to target putatively neutral loci, we used MAKER 2.31.8 to annotate the Cv_LA_1.0 assembly with training data from other galliforms and zebra finch (*Taeniopygia guttata*). Then, we used bedtools (20) to remove 7,335 SNPs that fell within regions annotated as genes, exons, five-prime-UTRs, or three-prime-UTRs in our reference genome (15). To avoid selecting linked sites, we used a custom Python script (21) to filter our dataset to include only one SNP per 100 kb window (22), which removed an additional 19,386 SNPs. Finally, we used

VCFtools (19) to remove 18 SNPs that fell on scaffolds <1 Mbp in our reference genome (N50 = 66.871 Mb). This produced a final dataset of 3,024 SNPs. We used bedtools to generate coordinates of the 120 bp sequence region surrounding each variable SNP in our reference genome, and we sent these coordinates and the draft genome assembly to Daicel Arbor Biosciences (Ann Arbor, MI, USA) for bait design. Arbor staff soft-masked simple and repetitive elements in each sequence using RepeatMasker (23) and replaced strings of ten or fewer unknown ("N") positions with a thymine. Because we were interested in enriching RAD loci from historical museum specimens with differing degrees of DNA degradation, Arbor staff designed three, 80 nt baits targeting each SNP where baits overlapped by 60 bp and spanned a 120 bp sequence region. Arbor staff then used NCBI BLAST (24) to compare 9,042 candidate bait sequences against themselves and our reference genome, removed non-specific bait sequences, and estimated the hybridization melting temperature ($T_m$) of all remaining baits. After discarding the BLAST hit with the highest $T_m$ for each bait candidate, we kept bait sequences with fewer than ten hits between 62.5 - 65° C and fewer than four hits above 65° C, which eliminated 1,266 bait sequences. Finally, we discarded 753 loci with fewer than three viable bait sequences, producing a final set of 7,053 baits targeting 2,351 SNPs. Daicel Arbor Biosciences synthesized this custom set of biotinylated RNA baits as a MYbaits Custom Kit.

**Library preparation, target enrichment, and sequencing**

For tissue samples, we followed the 3RAD library preparation protocol that we used to collect preliminary RAD data for bait design. We prepared three 3RAD pools, each containing eight uniquely dual-indexed individual libraries (pools included several samples not analyzed as part of this study). To identify and remove PCR duplicates prior to downstream analyses, we tagged each pool with a unique iTru7 index and a random octamer iTru5-8N index (4, 25). For toepad samples, we prepared standard dual-indexed genomic libraries of each using the KAPA Hyper Prep library preparation kit (F. Hoffmann-La Roche AG) and custom-indexes (26) at one-half volume. We amplified libraries using 12-16 cycles of PCR (98° C for 45 seconds; 12-16 cycles of 98° C for 15 seconds, 60° C for 30 seconds, 72° C for 30 seconds; 72° C for 1 minute) followed by 3X SpeedBead clean-up to remove unincorporated primers. Prior to enrichment, we quantified libraries using a Qubit Fluorometer (Life Technologies, Inc.) and combined toepad libraries into pools of eight using equimolar ratios so that specimens of similar age were mixed together.

We performed enrichment on all groups of eight libraries using our custom RADcap bait set following the myBaits Hybridization Capture for Targeted NGS manual v4.01, except that we substituted chicken C0t-1 (Applied Genetics Laboratories, Inc.) for Block C (human C0t-1) included in the myBaits kit. After enrichment, we performed 14-16 cycles of PCR recovery, and we cleaned the resulting reactions using the Qiagen GeneRead Size Selection Kit. We then ran post-enrichment pools on a Bioanalyzer (Agilent Technologies, Inc.) to verify peak size distributions and ensure the absence of adapter-dimers. Finally, we quantified enrichment pools using the KAPA qPCR quantification kit, and we combined pools at equimolar ratios prior to collecting sequence data using two lanes of 150-bp paired-end (PE150) sequencing on an Illumina NovaSeq 6000 (Novogene, Sacramento, CA).

**Preliminary RADcap data processing**

Because tissue and toepad libraries were prepared with different types of indexes, we initially processed each sample type separately. For the tissue samples, we used BBMap (27) to demultiplex sequencing lanes into pools using their unique iTru7 indexes. Then, we used *process_radtags* in Stacks v2.5 (14) to demultiplex each pool to individual samples, check resulting samples for the appropriate cut sites, and trim samples for adapters. Following adapter trimming, we removed PCR duplicates from each sample using the *clone_filter* tool in Stacks, which retains only a single read in instances when multiple reads contain identical iTru5-8N indexes and identical insert sequences. For the toepad samples, we used BBMap (27) to demultiplex each sample by their iTru7 and iTru5 indexes and illumiprocessor (28) to remove the indexes from the sequence data. The protocols we followed for processing our data are available on the Faircloth lab website (29, 30).

**SNP calling & filtering**

After preliminary processing of the RADcap data, we called SNPs using a parallel implementation of the Best Practices for Variant Discovery (31, 32). In brief, we used bwa v0.7.17 (16) and samtools v1.10 (17) to align demultiplexed and adapter-trimmed reads to the bobwhite reference genome (15). Following alignment, we used GATK v4.1.9.0 (33) to add read groups to each BAM file and mark and remove duplicates from the toepad samples. We did not mark duplicates for the tissue samples because we removed them during preliminary processing of the RADcap data. We indexed the final BAM files for all samples using samtools.

We performed an initial round of variant calling and filtering using a subset of our samples, from which we derived a set of “known variants" to use for base quality score recalibration across all samples (31). Because our dataset comprised fresh tissue samples and toepads from historical museum specimens, which often have higher levels of DNA damage and lower coverage than tissues (5), we selected 51 tissue samples (including the 14 tissue samples in this dataset and 37 additional *Colinus* samples not used as part of this study) for initial variant discovery. We called variants within 5bp of targeted SNPs across all samples and produced a single VCF file using HaplotypeCaller (-ERC GVCF), GenomicsDBImport (default parameters), and GenotypeGVCFs (default parameters). After calling variants, we used VCFtools v0.1.13 (19) to filter the resulting VCF to the 2,350 SNPs we targeted while removing indels, sites with less than 30x coverage, site quality and genotype quality scores below 30, non-biallelic sites, and sites with more than 50% missing data. This produced a VCF file containing 2,172 high-quality SNPs (92% of those targeted). Using this VCF file as our file of known variants, we performed base quality score recalibration across all sample BAM files (including those tissues from which the known variants were called) using BaseRecalibrator (default parameters) and ApplyBQSR (default parameters). We then called SNPs in all recalibrated BAMs using the same procedure as above, this time limiting the intervals in the genomic database to include the full 120 bp region surrounding our 2,350 targeted SNP loci.

Because the analyses we implemented assume sites are putatively neutral and unlinked (34, 35), we used VCFtools to filter SNPs in the resulting VCF file to the single locus targeted during bait design. We also used VCFtools to exclude: any loci determined to be indels, sites with less than a minimum depth of 10, sites with more than 10% missing data, and sites out of Hardy-Weinberg equilibrium due to excess heterozygosity, which can indicate genotyping errors (36). Because missing data can bias population structure assignment (37), we removed individuals missing more than 25% of calls at the remaining sites.

Then, because particular filters applied to a set of variable SNPs can affect different types of analyses in unexpected ways (38), we produced two types of VCF files: (Set1) for population genetic summary statistics, genetic structure, and phylogenetic analyses, and (Set2) for demographic analysis. To produce the first set of files (Set1), we used VCFtools to remove loci having a minor allele count (MAC) less than three (39), and we output files: (Set1a) including all bobwhite samples + the outgroup individuals (ingroup+outgroup); and (Set1b) including only the Cuban, USA, and Mexican populations of bobwhites (ingroup only). The inclusion of the outgroup individuals in this data file allowed us to test whether there was any allele sharing between Black-throated Bobwhites and Cuban bobwhites because some populations of Black-throated Bobwhites on the Yucatán Peninsula are physically closer to Cuba than populations of Northern Bobwhites in either Florida or the Caribbean slope of southern Mexico. To produce the second file for demographic analyses (Set2), we removed the MAC filtering because rare alleles can be important for inferring demographic history (40), and we included all bobwhites + the outgroup individuals so that we could include an external constraint on the common ancestor of the ingroup and outgroup (see below).

### Patterns of population genetic diversity

To assess population structure and potential admixture among populations, we calculated ancestry coefficients in sNMF (34) using both Set1a and Set1b VCF files. Specifically, we performed an initial set of analyses to determine the optimal K-value and the appropriate regularization parameter ($\alpha$) following the guidelines outlined by the program authors (34); for both datasets, we selected a regularization parameter of 100. After selecting these variables, we ran 100 analyses and selected the run with the lowest cross-entropy criterion score for each input file. To further characterize the patterns of genetic clustering in our dataset, we performed a discriminant analysis of principal components (DAPC) using VCFR v1.12.0 (41) and adegenet v2.1.3 (42) in R v4.0.3 (43). Following the guidelines outlined by the adegenet authors (44), we computed an initial set of principal components (PCs) to determine the optimal K-value, which we set as the number of populations for the DAPC. Because retaining too many PCs can lead to overfitting the data and erroneous population assignment (44), we retained the first 80 PCs, which explained 90% of the variance in the dataset, but we retained all eigenvectors because our dataset contained fewer than 10 genetic clusters. Finally, because both sNMF and DAPC require *a priori* selection of the number of clusters present in the data, we performed a principal components analysis (PCA) using SNPRelate (45) in R v4.0.3 (43).

To describe patterns of differentiation and genetic diversity within and among bobwhites (ingroup only), we calculated population genetic summary statistics (observed and expected heterozygosity, allelic richness, nucleotide diversity, inbreeding coefficients, and $F_{ST}$ between each pair of populations) using the Set1b VCF file. Specifically, we used hierfstat v0.5-7 (46) in R v4.0.3 (43) to calculate observed and expected heterozygosity, allelic richness, and inbreeding coefficients for each population and VCFtools (19) to calculate nucleotide diversity for each population, as well as $F_{ST}$ between each pair of populations. Finally, we calculated the number of private alleles in each population using a custom Python (47) script, which can be found on GitHub (48).

### Phylogenetic analyses

To investigate the evolutionary relationships among populations, we estimated species trees using SNAPP v1.5.1 (49) implemented in Beast v2.6.3 (50). We chose SNAPP because it was designed to model the coalescent process using allele frequencies from biallelic SNP data, and it produces output that enables users to visualize conflicting signals due to allele sharing among multiple populations, which we suspected in our dataset based on the history of multiple introductions into Cuba. However, SNAPP is computationally intensive, which precluded us from estimating a species tree from the entire dataset.

To work around this limitation, we used two different types of subsampling schemes. In the first scheme, we were interested in inferring the relationships and extent of allele sharing among individuals in each of the four populations identified by our sNMF analyses (Cuba, northern Mexico, Southern Mexico, and USA) (Table S7), so we used R to create five replicate subsamples containing five randomly sampled individuals from each of these four populations (i.e., 20 individuals in each of five subsamples). In the second scheme, we were more interested in trying to understand the complicated history of introductions across Cuba and the degree of allele sharing between mainland populations and individuals sampled at each of the five, distinct Cuban localities. So, we used R to create locality-specific data subsets. Each subset included all individuals from a distinct sampling locality in Cuba (3-6 individuals per locality) and two randomly selected individuals from each of the twelve mainland subpopulations (three subpopulations in USA, three subpopulations in northern Mexico, and six subpopulations in Southern Mexico) (Table S1). This resulted in a total of 27-30 individuals in each of five analyses (Table S7).

Once we selected the individuals in subsamples for both subsampling schemes, we used VCFtools to output VCF files containing SNP data for each by extracting data from the Set1b VCF file, we converted each VCF file to SNAPP format using VCF2SNAPP (51) in R v4.0.3 (43), and we used

BEAUti v2.6.3 (50) to prepare subset-specific input files for reach run. We ran each analysis in SNAPP with default parameters for 2 million iterations, sampling every 1,000 iterations and discarding the first 10% of sampled iterations as burn-in. We used Tracer v1.7.2 (52) to visualize the traces of estimated parameters and to ensure that effective sample size of each (ESS) was greater than 200 (53). Finally, we visualized the resulting species trees and posterior distribution of gene trees using DensiTree v2.2.7 (54, 55).

**Demographic analyses**

Inferring evolutionary relationships and visualizing allele sharing among populations provide one mechanism for understanding population origins, but these types of analyses do not typically allow objective comparisons of alternative evolutionary scenarios, including comparisons of the timing of evolutionary events and/or the source(s) and direction of migrants to specific populations. This is particularly true in the absence of suitable fossil calibration points, which is currently the case for Cuban bobwhites (56), and over the recent timescales used in two of the hypotheses explaining the origin of Cuban bobwhites: both D'Orbigny and Parkes suggested that bobwhites were not present on Cuba until introduced by Europeans during the last 500 years. To perform these types of analyses, we used momi2 (35) with the site frequency spectrum (SFS) derived from the Set2 VCF file.

Specifically, we tested two categories of models given our data: 1) single source models, in which one of the mainland populations (the ancestor of Northern + Southern Mexico, northern Mexico, Southern Mexico, or USA) founded the Cuban population, followed by a single pulse of migration from the same founding population to the Cuban population; and 2) multiple source models, in which one of the mainland populations founded the Cuban population, followed by a single pulse of migration from a different mainland population during the last 500 years – a design consistent with the recorded history of bobwhite introductions to Cuba since the arrival of Europeans (57, 58). For models in the multi-source category, we constrained each to always include the USA, either as the founding population or as the source of the more recent pulse of migration to Cuba because SNAPP analyses always showed allele sharing between the USA and Cuba. For example, we did not model a scenario in which northern Mexico was the founder of the Cuban population, followed by a recent pulse of migration from southern Mexico because this was inconsistent with the SNAPP results. To allow tests of hypotheses regarding the timing of bobwhite arrival on Cuba, we incorporated five different temporal scenarios to all models in both categories for the founding of the Cuban population – specifically, that the Cuban bobwhite population was founded: 1) within 0.5 kya (since European arrival on Cuba; (59); 2) between 0.5-5.0 kya (since Indigenous arrival on Cuba; (60)); 3) between 5.0-11.7 kya (since the Pleistocene-Holocene Transition; (61)); 4) between 11.7-23.0 kya (since the last glacial maximum; (61)); or 5) between 23.0-140.0 kya (since the penultimate glacial maximum; (62). In total, this produced 45 separate models (Supplementary Figure S2; Supplementary Table S9).

Before running momi2, we set the effective population size of the root to $3.35 \times 10^5$, the mean value estimated across multiple G-PhoCs runs for the population ancestral to Northern Bobwhites and their sister species, Black-throated Bobwhites (see below); we specified a generation time of 1.22 years, which was the median value estimated from multiple radiotelemetry and survivorship studies of wild bobwhites (63); and we specified a mutation rate of $1.91 \times 10^{-9}$ sites per year, which was estimated from chickens (64). Finally, we constrained the divergence time between Northern Bobwhites and Black-throated Bobwhites to have occurred since 1.563 Ma, which was the older bound of the 95% highest posterior density interval estimated for their divergence in a time-calibrated phylogeny (65).

To perform an initial comparison of models in a computationally tractable way, we ran each of the 45 models 10 times and computed corrected Akaike information criterion (AICc) scores (66, 67) for all models in each set of the 10 model runs. We then used AIC-based model comparison (68) to rank and compare models (Table S10). Only three models fell within the confidence set (69) among any of the 10 runs. To ensure that we sufficiently searched the likelihood surface for each of the three models in the confidence set, we completed an additional 90 runs of each, selected the best (highest) log-likelihood value for each model out of 100 (10+90) runs, and used the best log-likelihood obtained for each of the three top models to compute final AICc scores, delta AICc values, and Akaike weights following Burnham and Anderson (68).

**Demographic parameter estimates with G-PhoCs**

We used G-PhoCs (70) to estimate the effective population size of the common ancestor of Northern Bobwhites and their sister species, Black-throated Bobwhites, which is one of the input parameters required for demographic modeling in momi2. Because G-PhoCs requires alignments of complete loci (including invariant sites) as input, we used VCFtools (19) to filter the VCF file we produced by calling SNPs within the full 121 bp region surrounding our 2,350 targeted SNP loci (the starting file from which we produced both Set1 and Set2 VCF files). To produce a dataset for use with G-PhoCs containing all 109 individuals (ingroup + outgroup), we used VCFtools to retain all SNPs within each locus (not just the single targeted SNP) and to exclude: any loci determined to be indels, sites with less than a minimum depth of 10, and sites with more than 10% missing data. With this VCF file, we used BCFTools consensus v1.12 (71) to apply the variants from each sample to the bobwhite reference genome sequence (15) to create a fasta file containing the complete 121 bp sequences of all 2,350 loci for each sample (e.g. bcftools consensus -f <bobwhite_ref_genome.fasta> -H I -I -s <sample_name> -o <sample_name_consensus.fasta> <filtered_VCF_file>). We combined the resulting fasta files for all samples in bash and used the Phyluce script phyluce_assembly_explode_get_fastas_file (72) to output a single fasta file for each locus. We used mafft (-no-trim, -ambiguous, -incomplete-matrix) implemented in phyluce_align_seqcap_align to produce relaxed-phylip format alignments of all loci, which we concatenated to produce the G-PhoCs input file containing 2,350 loci (121 bp each) for all 109 taxa.

Due to the computational constraints of G-PhoCs, we could not include all individuals in a single analysis, so we instead took an approach similar to our SNAPP analyses and used R to randomly sample two non-overlapping sets of four individuals from each of the four geographic populations identified in the sNMF analysis (Cuba, USA, northern Mexico, and southern Mexico), as well as the outgroup, Black-throated Bobwhites (20 individuals total). We used population divergence models allowing bidirectional gene flow between pairs of modern, geographically adjacent mainland populations (e.g. between USA and northern Mexico, but not between USA and southern Mexico), as well as between the Cuban population and all modern and historical (e.g. the common ancestor of northern and southern Mexico) mainland populations, including the outrgroup (migration model). We used the standard settings for the MCMC described by Gronau et al. (70) and Freedman et al. (73), including diffuse gamma prior distributions for all demographic parameters. The shape ($\alpha$) and scale ($\beta$) parameters that define the gamma prior distributions for each of the demographic parameters were as follows: $\alpha$ = 1 and $\beta$ = 10,000 for both divergence time ($\tau$) and population size ($\theta$), and $\alpha$ = 0.002 and $\beta$ = 0.00001 for the migration rates ($m$). We conducted initial MCMC runs for each analysis with auto-fine-tuning to evaluate run speed, convergence, and mixing, before conducting final analyses using 100,000-200,000 generations of burn-in, followed by 800,000-900,000 sampled generations. We evaluated the convergence of final runs using Tracer v1.7.2 (52).

Because our randomly-sampled SNAPP trees resolved two different topologies with respect to the placement of the Cuban population (sister to either USA or both Mexican populations), we conducted two separate runs for both randomly-sampled datasets: a run specifying the Cuba + USA topology as the underlying guide tree and a run specifying the Cuba + Mexico topology as the underlying guide tree. In total, we ran four analyses in G-Phocs.

We converted the posterior distributions for the estimates of different parameters from mutation scale to generations and individuals by assuming an average mutation rate of 1.91 x $10^{-9}$ sites per year, which was estimated from chickens (64), and a generation time of 1.22 years, which was the median value estimated from multiple radiotelemetry and survivorship studies of wild bobwhites (63). Because all four runs produced similar values for the effective population size of the common ancestor of Northern and Black-throated Bobwhites, we used the mean value across all runs (3.35 x $10^5$, 95% CI 2.82 x $10^4$) as the input parameter for our momi2 analyses.

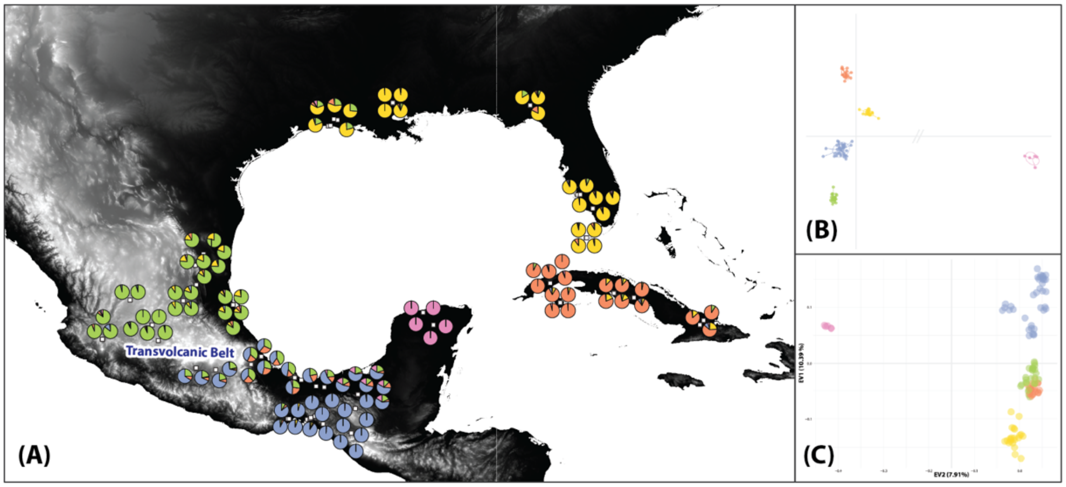


**Fig. S1. Population structure of Northern and Black-throated Bobwhites from Cuba, the USA, and Mexico. (A)** sNMF results for best-fit K-value of five populations. Pie charts show admixture proportions for each individual plotted by sampling locality (points have been jittered around exact coordinates to allow easier viewing). **(B)** DAPC results for best-fit five population clusters: USA (yellow), Northern Mexico (green), Southern Mexico (blue), Cuba (orange), and Black-throated Bobwhites (pink). (C) PCA results for the same set of individuals. Clusters have been colored as in (A-B) to reflect population assignment from sNMF and PCA.

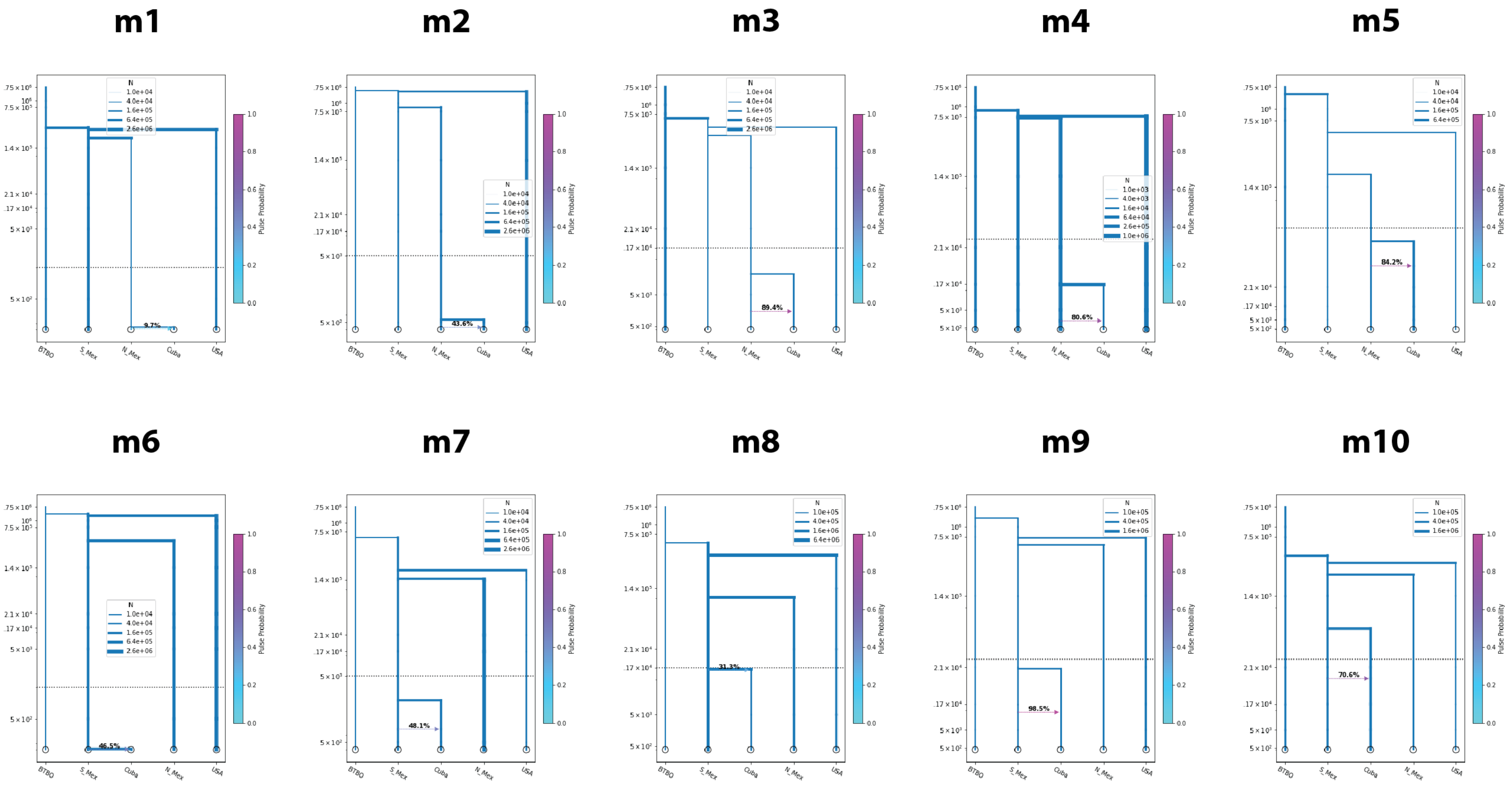


**Fig. S2. Schematics of the 45 demographic models tested with momi2**. Parameter values are randomly generated within the provided constraints (see Materials and Methods) and **do not represent real output**. m1-m20 are single source models; m21-m45 are multiple source models. BTBO = Black-throated Bobwhites; S_Mex = southern Mexico; N_Mex = northern Mexico. For descriptions of each model, see Supplementary Table S9.
*(figure cont'd.)*

# m11

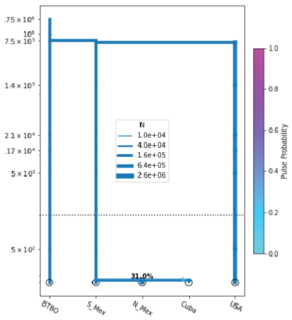


# m12

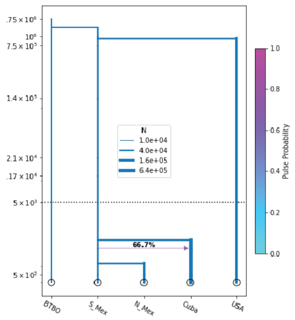


# m13

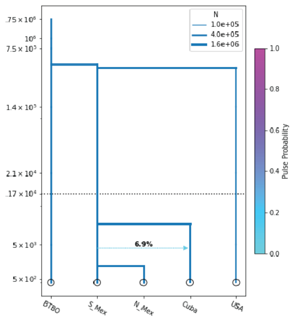


# m14

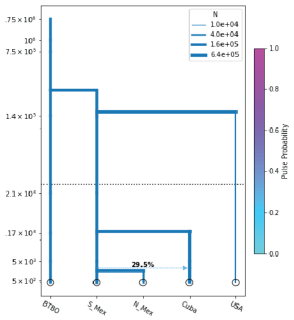


# m15

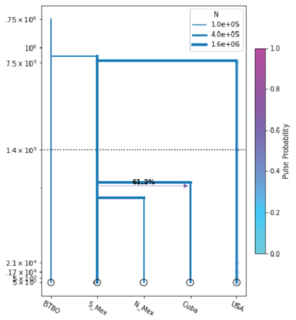


# m16

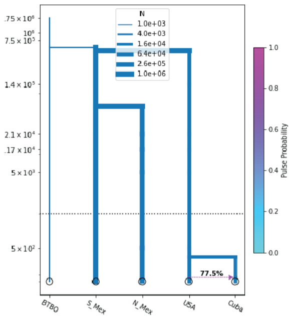


# m17

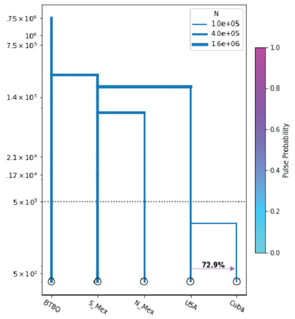


# m18

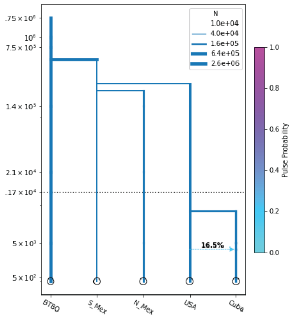


# m19

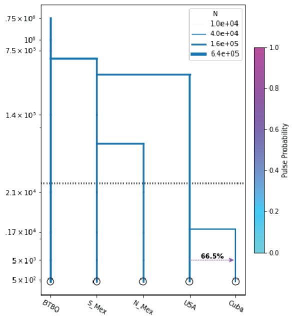


# m20

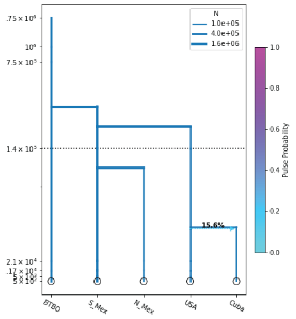

**m21**

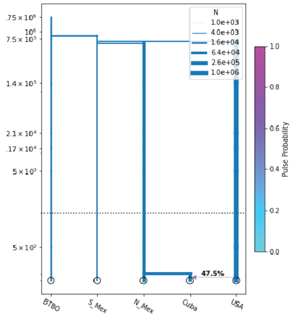


**m22**

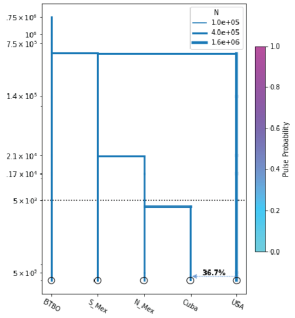


**m23**

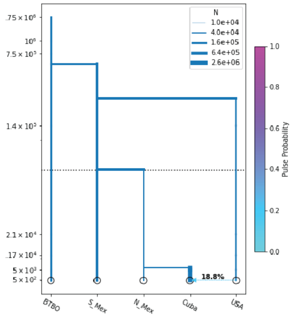


**m24**

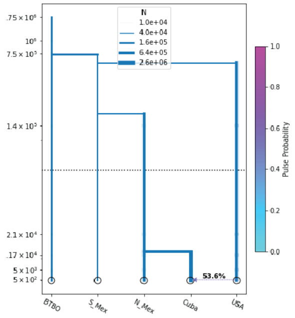


**m25**

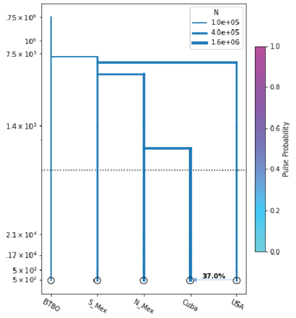


**m26**

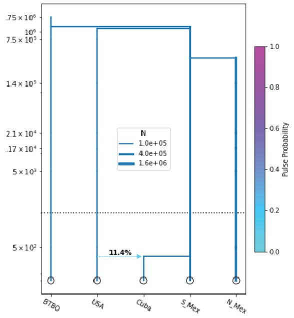


**m27**

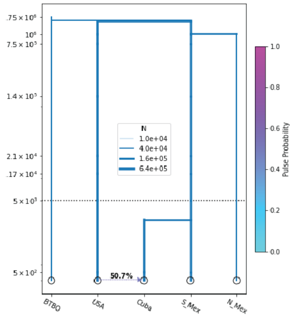


**m28**

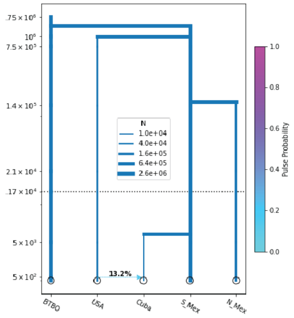


**m29**

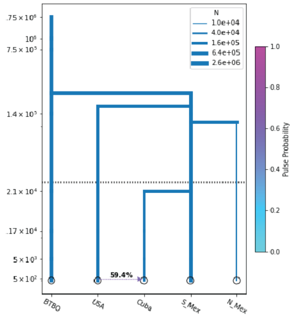


**m30**

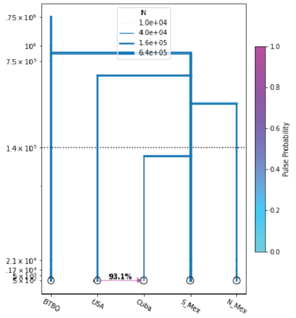

## m31

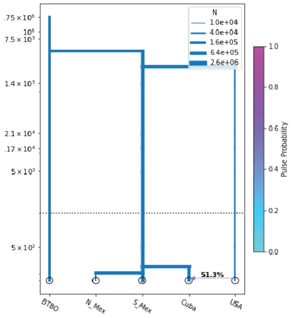


## m32

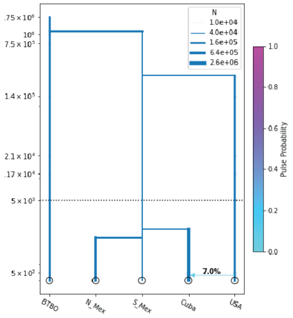


## m33

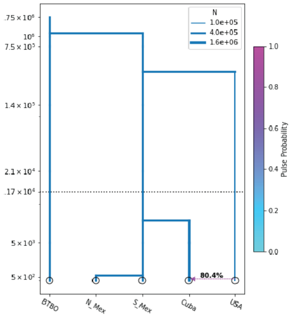


## m34

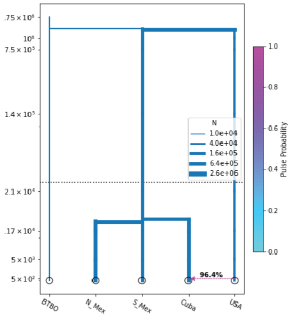


## m35

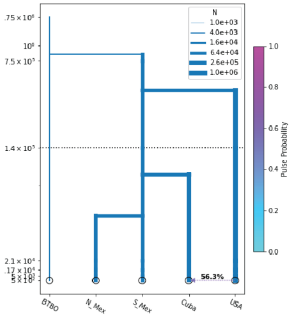


## m36

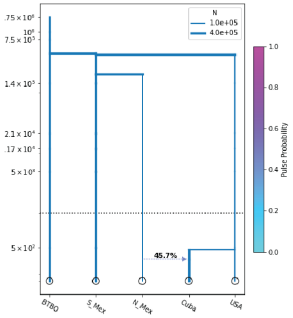


## m37

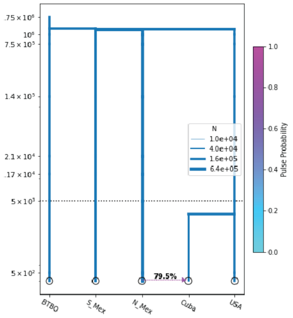


## m38

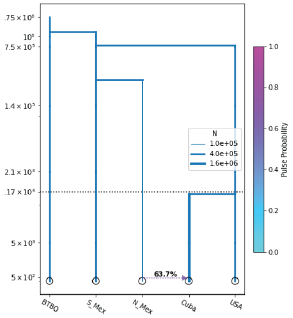


## m39

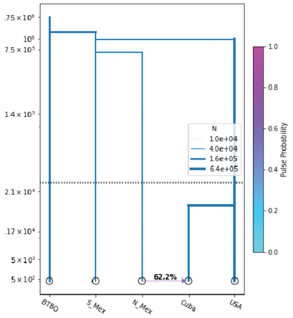


## m40

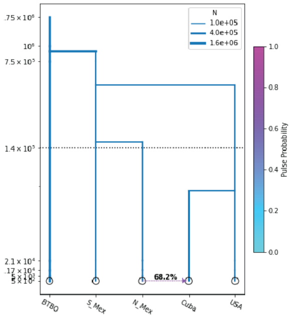

**m41**

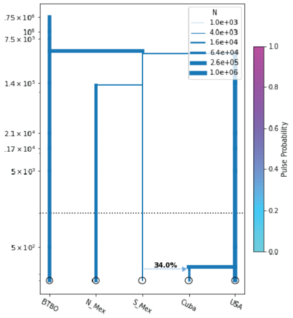


**m42**

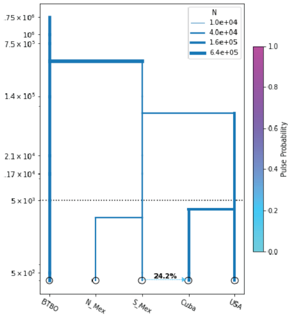


**m43**

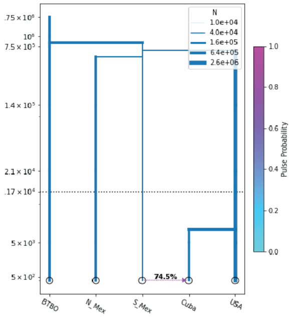


**m44**

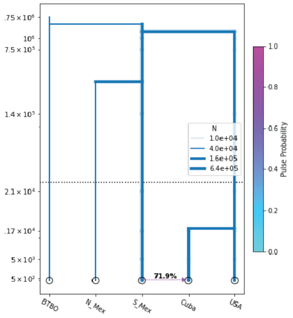


**m45**

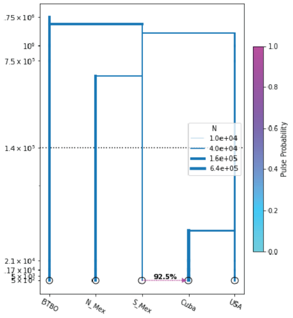

**Table S1.** Sample information and sequencing summary statistics. Spp. = Species; Mus. = Museum; Cat. No. = Catalog or Accession Number; Coll. Year = Collection Year; Avg. Depth of Cov. = Average Depth of Coverage. *C. n.* = *Colinus nigrogularis* (Black-throated Bobwhites); *C. v.* = *Colinus virginianus* (Northern Bobwhites). Subspecies identifications are taken from the collection where the specimen is accessioned and may have been assigned by locality without detailed diagnosis. The number of SNPs for each sample in the Set1a population structure dataset is shown.

| Museum | Cat. No | Species | Subspecies / Subpopulation | Locality | Sample Type | Coll. Year | Raw Read Pairs | Avg. Depth of Cov. | SNPs |
|---|---|---|---|---|---|---|---|---|---|
| YPM | 14442 | *C. nigrogularis* | *caboti* | Yucatán, Mexico | toepad | 1949 | 2623428 | 123.59 | 1267 |
| YPM | 14443 | *C. nigrogularis* | *caboti* | Yucatán, Mexico | toepad | 1949 | 1629449 | 90.277 | 1267 |
| YPM | 14424 | *C. nigrogularis* | *persiccus* | Yucatán, Mexico | toepad | 1950 | 1183498 | 70.1351 | 1267 |
| YPM | 14431 | *C. nigrogularis* | *persiccus* | Yucatán, Mexico | toepad | 1950 | 1224125 | 77.1922 | 1267 |
| YPM | 14433 | *C. nigrogularis* | *persiccus* | Yucatán, Mexico | toepad | 1950 | 1184843 | 69.6433 | 1267 |
| MCZ | 280778 | *C. virginianus* | *cubanensis x floridanus* | Cienfuegos Province, Cuba | toepad | 1941 | 1183188 | 53.0269 | 1267 |
| MCZ | 280779 | *C. virginianus* | *cubanensis x floridanus* | Cienfuegos Province, Cuba | toepad | 1941 | 1236032 | 53.2421 | 1267 |
| MCZ | 280780 | *C. virginianus* | *cubanensis x floridanus* | Cienfuegos Province, Cuba | toepad | 1941 | 1339663 | 51.9715 | 1267 |
| MCZ | 280781 | *C. virginianus* | *cubanensis x floridanus* | Cienfuegos Province, Cuba | toepad | 1941 | 1966706 | 73.5191 | 1267 |
| MCZ | 114910 | *C. virginianus* | *cubanensis* | Holguín Provice, Cuba | toepad | 1904 | 676213 | 19.2356 | 1081 |
| MCZ | 114911 | *C. virginianus* | *cubanensis* | Holguín Provice, Cuba | toepad | 1904 | 1044236 | 32.6224 | 1242 |
| MCZ | 114912 | *C. virginianus* | *cubanensis* | Holguín Provice, Cuba | toepad | 1904 | 774510 | 27.076 | 1210 |
| YPM | 33310 | *C. virginianus* | *cubanensis* | Isla de la Juventud, Cuba | toepad | 1955 | 2473649 | 127.606 | 1267 |
| YPM | 33311 | *C. virginianus* | *cubanensis* | Isla de la Juventud, Cuba | toepad | 1955 | 1150259 | 76.6377 | 1267 |
| YPM | 33312 | *C. virginianus* | *cubanensis* | Isla de la Juventud, Cuba | toepad | 1955 | 1025973 | 68.8102 | 1267 |
| YPM | 33313 | *C. virginianus* | *cubanensis* | Isla de la Juventud, Cuba | toepad | 1955 | 660108 | 32.6164 | 1259 |
| YPM | 33315 | *C. virginianus* | *cubanensis* | Pinar del Río Province, Cuba | toepad | 1955 | 999915 | 64.6196 | 1267 |
| YPM | 33316 | *C. virginianus* | *cubanensis* | Pinar del Río Province, Cuba | toepad | 1955 | 544333 | 47.5493 | 1267 |
| YPM | 33317 | *C. virginianus* | *cubanensis* | Pinar del Río Province, Cuba | toepad | 1955 | 1272346 | 81.0012 | 1267 |
| YPM | 33319 | *C. virginianus* | *cubanensis* | Pinar del Río Province, Cuba | toepad | 1955 | 588519 | 51.7583 | 1267 |

| Museum | Cat. No | Species | Subspecies / Subpopulation | Locality | Sample Type | Coll. Year | Raw Read Pairs | Avg. Depth of Cov. | SNPs |
|---|---|---|---|---|---|---|---|---|---|
| YPM | 33320 | *C. virginianus* | *cubanensis* | Pinar del Río Province, Cuba | toepad | 1955 | 528806 | 40.8834 | 1266 |
| YPM | 58010 | *C. virginianus* | *cubanensis* | Sancti Spíritus Province, Cuba | toepad | 1925 | 1229338 | 58.349 | 1267 |
| YPM | 58011 | *C. virginianus* | *cubanensis* | Sancti Spíritus Province, Cuba | toepad | 1925 | 1433390 | 67.698 | 1267 |
| YPM | 58012 | *C. virginianus* | *cubanensis* | Sancti Spíritus Province, Cuba | toepad | 1925 | 1190771 | 63.3575 | 1267 |
| MLZ | 39388 | *C. virginianus* | *graysoni* | Aguascalientes, Mexico | toepad | 1944 | 932065 | 58.8235 | 1266 |
| MLZ | 39389 | *C. virginianus* | *graysoni* | Aguascalientes, Mexico | toepad | 1944 | 2207436 | 81.7527 | 1267 |
| MLZ | 27065 | *C. virginianus* | *coyolcos* | Chiapas, Mexico | toepad | 1940 | 1458879 | 60.8448 | 1266 |
| WFVZ | 4145 | *C. virginianus* | *coyolcos* | Chiapas, Mexico | toepad | 1957 | 1010022 | 63.1407 | 1267 |
| WFVZ | 4146 | *C. virginianus* | *coyolcos* | Chiapas, Mexico | toepad | 1957 | 1605974 | 91.1444 | 1267 |
| WFVZ | 4147 | *C. virginianus* | *coyolcos* | Chiapas, Mexico | toepad | 1957 | 502161 | 45.817 | 1267 |
| WFVZ | 17806 | *C. virginianus* | *insignis* | Chiapas, Mexico | toepad | 1941 | 2023490 | 95.2276 | 1267 |
| WFVZ | 17807 | *C. virginianus* | *insignis* | Chiapas, Mexico | toepad | 1945 | 638590 | 45.6562 | 1263 |
| WFVZ | 17808 | *C. virginianus* | *insignis* | Chiapas, Mexico | toepad | 1941 | 1561233 | 77.8287 | 1267 |
| LACM | 24445 | *C. virginianus* | *insignis* | Chiapas, Mexico | toepad | 1954 | 932354 | 68.3659 | 1267 |
| WFVZ | 3680 | *C. virginianus* | *insignis* | Chiapas, Mexico | toepad | 1957 | 2017663 | 105.631 | 1266 |
| WFVZ | 8854 | *C. virginianus* | *insignis* | Chiapas, Mexico | toepad | 1962 | 1067847 | 56.0237 | 1266 |
| WFVZ | 8856 | *C. virginianus* | *insignis* | Chiapas, Mexico | toepad | 1962 | 2282134 | 79.737 | 1266 |
| WFVZ | 8857 | *C. virginianus* | *insignis* | Chiapas, Mexico | toepad | 1962 | 999816 | 50.6064 | 1267 |
| WFVZ | 8858 | *C. virginianus* | *insignis* | Chiapas, Mexico | toepad | 1962 | 917275 | 54.1737 | 1267 |
| UMMZ | 104504 | *C. virginianus* | *minor* | Chiapas, Mexico | toepad | 1939 | 1319881 | 67.6755 | 1266 |
| UMMZ | 104507 | *C. virginianus* | *minor* | Chiapas, Mexico | toepad | 1939 | 1428645 | 70.7463 | 1266 |
| UMMZ | 104508 | *C. virginianus* | *minor* | Chiapas, Mexico | toepad | 1939 | 3244564 | 114.568 | 1266 |
| MLZ | 43826 | *C. virginianus* | *minor* | Chiapas, Mexico | toepad | 1946 | 1164625 | 65.4524 | 1266 |

| Museum | Cat. No | Species | Subspecies / Subpopulation | Locality | Sample Type | Coll. Year | Raw Read Pairs | Avg. Depth of Cov. | SNPs |
|---|---|---|---|---|---|---|---|---|---|
| MLZ | 43827 | *C. virginianus* | *minor* | Chiapas, Mexico | toepad | 1946 | 927859 | 49.9582 | 1267 |
| MLZ | 44393 | *C. virginianus* | *minor* | Chiapas, Mexico | toepad | 1946 | 1098147 | 60.6643 | 1264 |
| MLZ | 35576 | *C. virginianus* | *graysoni* | Guanajuato, Mexico | toepad | 1943 | 911234 | 55.0848 | 1266 |
| MLZ | 36640 | *C. virginianus* | *graysoni* | Guanajuato, Mexico | toepad | 1943 | 1457618 | 77.6506 | 1267 |
| MLZ | 36726 | *C. virginianus* | *graysoni* | Guanajuato, Mexico | toepad | 1943 | 1386455 | 73.5428 | 1267 |
| MLZ | 25606 | *C. virginianus* | *graysoni* | Jalisco, Mexico | toepad | 1940 | 2094447 | 72.4753 | 1266 |
| MLZ | 25607 | *C. virginianus* | *graysoni* | Jalisco, Mexico | toepad | 1940 | 960556 | 54.0426 | 1267 |
| MLZ | 25609 | *C. virginianus* | *graysoni* | Jalisco, Mexico | toepad | 1940 | 835999 | 56.889 | 1267 |
| MLZ | 57882 | *C. virginianus* | *graysoni* | Michoacan, Mexico | toepad | 1954 | 1204139 | 72.3225 | 1267 |
| MLZ | 57884 | *C. virginianus* | *graysoni* | Michoacan, Mexico | toepad | 1954 | 806265 | 62.4797 | 1267 |
| SDNHM | 54282 | *C. virginianus* | *nigripectus* | Morelos, Mexico | tissue | 2014 | 1970606 | 49.2014 | 1255 |
| SDNHM | 54283 | *C. virginianus* | *nigripectus* | Morelos, Mexico | tissue | 2014 | 1687059 | 48.3144 | 1255 |
| SDNHM | 54284 | *C. virginianus* | *nigripectus* | Morelos, Mexico | tissue | 2014 | 1175291 | 46.9292 | 1250 |
| WFVZ | 12490 | *C. virginianus* | *coyolcos* | Oaxaca, Mexico | toepad | 1964 | 5980941 | 208.844 | 1267 |
| WFVZ | 16981 | *C. virginianus* | *coyolcos* | Oaxaca, Mexico | toepad | 1966 | 507482 | 40.0905 | 1265 |
| WFVZ | 20968 | *C. virginianus* | *coyolcos* | Oaxaca, Mexico | toepad | 1964 | 2576458 | 101.023 | 1266 |
| WFVZ | 20996 | *C. virginianus* | *coyolcos* | Oaxaca, Mexico | toepad | 1964 | 3390687 | 132.319 | 1267 |
| WFVZ | 48912 | *C. virginianus* | *coyolcos* | Oaxaca, Mexico | toepad | 1964 | 2851364 | 103.946 | 1266 |
| WFVZ | 3679 | *C. virginianus* | *nigripectus* | Puebla, Mexico | toepad | 1957 | 1167505 | 74.39 | 1267 |
| MLZ | 53450 | *C. virginianus* | *graysoni* | San Luis Potosí, Mexico | toepad | 1952 | 1401199 | 78.963 | 1267 |
| MLZ | 53452 | *C. virginianus* | *graysoni* | San Luis Potosí, Mexico | toepad | 1952 | 2552249 | 138.505 | 1267 |
| MLZ | 53453 | *C. virginianus* | *graysoni* | San Luis Potosí, Mexico | toepad | 1952 | 1139027 | 79.4459 | 1267 |
| MLZ | 53454 | *C. virginianus* | *graysoni* | San Luis Potosí, Mexico | toepad | 1952 | 1715256 | 91.9453 | 1267 |

| Museum | Cat. No | Species | Subspecies / Subpopulation | Locality | Sample Type | Coll. Year | Raw Read Pairs | Avg. Depth of Cov. | SNPs |
|---|---|---|---|---|---|---|---|---|---|
| MLZ | 40200 | *C. virginianus* | *maculatus* | San Luis Potosí, Mexico | toepad | 1944 | 802006 | 54.1974 | 1267 |
| LSU | 27130 | *C. virginianus* | *godmani* | Tabasco, Mexico | toepad | 1961 | 1007796 | 46.3542 | 1267 |
| LSU | 27131 | *C. virginianus* | *godmani* | Tabasco, Mexico | toepad | 1961 | 560149 | 28.3856 | 1215 |
| MLZ | 45547 | *C. virginianus* | *aridus* | Tamaulipas, Mexico | toepad | 1947 | 1807526 | 93.9783 | 1267 |
| MLZ | 45549 | *C. virginianus* | *aridus* | Tamaulipas, Mexico | toepad | 1947 | 1293963 | 66.082 | 1267 |
| MLZ | 45554 | *C. virginianus* | *aridus* | Tamaulipas, Mexico | toepad | 1947 | 1435714 | 74.4548 | 1267 |
| LSU | 7700 | *C. virginianus* | *aridus* | Tamaulipas, Mexico | toepad | 1943 | 348256 | 28.4524 | 1216 |
| LSU | 7701 | *C. virginianus* | *aridus* | Tamaulipas, Mexico | toepad | 1943 | 269012 | 22.3961 | 1158 |
| LSU | 7702 | *C. virginianus* | *aridus* | Tamaulipas, Mexico | toepad | 1943 | 459450 | 30.2807 | 1225 |
| LSU | 28759 | *C. virginianus* | *godmani* | Veracruz, Mexico | toepad | 1962 | 844559 | 46.0056 | 1266 |
| LSU | 61133 | *C. virginianus* | *godmani* | Veracruz, Mexico | toepad | 1961 | 1215848 | 54.3092 | 1266 |
| MLZ | 39437 | *C. virginianus* | *maculatus* | Veracruz, Mexico | toepad | 1944 | 1733802 | 81.9646 | 1267 |
| MLZ | 39457 | *C. virginianus* | *maculatus* | Veracruz, Mexico | toepad | 1944 | 1388633 | 75.4528 | 1267 |
| MLZ | 40230 | *C. virginianus* | *maculatus* | Veracruz, Mexico | toepad | 1944 | 2299970 | 94.1391 | 1266 |
| MLZ | 40231 | *C. virginianus* | *maculatus* | Veracruz, Mexico | toepad | 1944 | 1421821 | 76.2831 | 1267 |
| MLZ | 40232 | *C. virginianus* | *maculatus* | Veracruz, Mexico | toepad | 1944 | 1581966 | 76.7021 | 1267 |
| LSU | 2828 | *C. virginianus* | *pectoralis* | Veracruz, Mexico | toepad | 1938 | 711919 | 40.4057 | 1262 |
| MLZ | 35866 | *C. virginianus* | *pectoralis* | Veracruz, Mexico | toepad | 1943 | 1303081 | 70.4869 | 1267 |
| MLZ | 35867 | *C. virginianus* | *pectoralis* | Veracruz, Mexico | toepad | 1943 | 2202699 | 83.4391 | 1267 |
| MLZ | 46867 | *C. virginianus* | *pectoralis* | Veracruz, Mexico | toepad | 1947 | 1213105 | 61.197 | 1267 |
| MLZ | 46883 | *C. virginianus* | *pectoralis* | Veracruz, Mexico | toepad | 1947 | 1015004 | 63.4616 | 1267 |
| MCZ | 14153 | *C. virginianus* | *floridanus* | Florida, USA | toepad | unknown | 1610026 | 51.4483 | 1266 |
| MCZ | 14154 | *C. virginianus* | *floridanus* | Florida, USA | toepad | unknown | 824293 | 32.3651 | 1251 |

| Museum | Cat. No | Species | Subspecies / Subpopulation | Locality | Sample Type | Coll. Year | Raw Read Pairs | Avg. Depth of Cov. | SNPs |
|---|---|---|---|---|---|---|---|---|---|
| MCZ | 14155 | *C. virginianus* | *floridanus* | Florida, USA | toepad | unknown | 1210297 | 41.7435 | 1264 |
| MCZ | 14156 | *C. virginianus* | *floridanus* | Florida, USA | toepad | unknown | 941645 | 31.1267 | 1240 |
| MCZ | 270594 | *C. virginianus* | *floridanus* | Florida, USA | toepad | 1941 | 978607 | 48.152 | 1266 |
| MCZ | 270595 | *C. virginianus* | *floridanus* | Florida, USA | toepad | 1941 | 1129491 | 55.7435 | 1267 |
| MCZ | 270596 | *C. virginianus* | *floridanus* | Florida, USA | toepad | 1941 | 1019351 | 56.8581 | 1267 |
| MCZ | 270597 | *C. virginianus* | *floridanus* | Florida, USA | toepad | 1941 | 1100614 | 35.2417 | 1252 |
| LSU | 48982 | *C. virginianus* | *floridanus* | Florida, USA | tissue | 2002 | 2368779 | 48.5943 | 1208 |
| LSU | 69842 | *C. virginianus* | *floridanus* | Florida, USA | tissue | 2006 | 1415605 | 71.6112 | 1267 |
| DMNH | 54851 | *C. virginianus* | *virginianus* | Georgia, USA | toepad | 1952 | 320405 | 22.3466 | 1183 |
| DMNH | 54852 | *C. virginianus* | *virginianus* | Georgia, USA | toepad | 1952 | 935289 | 47.4154 | 1265 |
| DMNH | 54854 | *C. virginianus* | *virginianus* | Georgia, USA | toepad | 1952 | 728856 | 45.5641 | 1265 |
| LSU | B-27329 | *C. virginianus* | *mexicanus* | Louisiana, USA | tissue | 1996 | 1436846 | 45.189 | 1237 |
| LSU | B-3283 | *C. virginianus* | *mexicanus* | Louisiana, USA | tissue | 1985 | 1783468 | 45.7386 | 1237 |
| LSU | B-3713 | *C. virginianus* | *mexicanus* | Louisiana, USA | tissue | 1986 | 1886602 | 45.8649 | 1229 |
| LSU | B-41222 | *C. virginianus* | *mexicanus* | Louisiana, USA | tissue | 2000 | 206636 | 34.8267 | 1011 |
| LSU | B-64490 | *C. virginianus* | *mexicanus* | Louisiana, USA | tissue | 2002 | 220077 | 35.8058 | 1021 |
| LSU | B-91876 | *C. virginianus* | *mexicanus* | Louisiana, USA | tissue | 2016 | 2483216 | 47.0595 | 1240 |
| LSU | B-91918 | *C. virginianus* | *mexicanus* | Louisiana, USA | tissue | 2017 | 1575122 | 48.3446 | 1255 |
| LSU | B-91919 | *C. virginianus* | *mexicanus* | Louisiana, USA | tissue | 2017 | 2473838 | 47.3422 | 1246 |
| LSU | B-91920 | *C. virginianus* | *mexicanus* | Louisiana, USA | tissue | 2017 | 2126906 | 46.2445 | 1242 |

**Table S2.** Sequencing summary statistics for tissues and toepads.

| Sample type | No. of samples | Mean raw reads ± 95% CI | Reads on target (%) | Duplicate reads (%) | Mean depth of coverage ± 95% CI |
|---|---|---|---|---|---|
| Tissues | 14 | 1,629,289 ± 377,407 | 95.7 | 4.96 | 47 ± 4 |
| Toepads | 95 | 1,338,505 ± 158,242 | 24.1 | 24.8 | 66 ± 6 |

**Table S3.** Population genetic summary statistics. For each population, the table shows the number of individuals, observed ($H_O$) and expected ($H_S$) heterozygosity, allelic richness, nucleotide diversity (**π**), and inbreeding coefficient ($F_{IS}$). All statistics were calculated Set1b dataset. N. Mex = northern Mexico; S. Mex = southern Mexico.

| Population | Sample size | $H_O$ | $H_S$ | Allelic richness | π | $F_{IS}$ |
|---|---|---|---|---|---|---|
| Cuba | 19 | 0.223 | 0.237 | 1.679 | 0.236 | 0.050 |
| N. Mex | 26 | 0.232 | 0.249 | 1.785 | 0.249 | 0.052 |
| S. Mex | 37 | 0.174 | 0.214 | 1.693 | 0.213 | 0.155 |
| USA | 22 | 0.260 | 0.281 | 1.893 | 0.280 | 0.062 |

**Table S4.** $F_{ST}$ between population pairs (lower half in white). Private alleles in black diagonal. Private alleles within pairs of populations in gray. N. Mex = northern Mexico; S. Mex = southern Mexico.

| **Population** | Cuba | N. Mexico | S. Mexico | USA |
|---|---|---|---|---|
| Cuba | 7 | 1 | 11 | 47 |
| N. Mex | 0.155 | 6 | 57 | 74 |
| S. Mex | 0.184 | 0.167 | 24 | 24 |
| USA | 0.126 | 0.135 | 0.226 | 59 |

**Table S5. Parameter estimates for the two best-supported demographic models.** Divergence ages are given in years. $N_e$ = effective population size.

| Parameter | Southern Mexico founder (0.5-5 kya), subsequent migration from USA (<0.5 kya) | Southern Mexico founder (<0.5 kya), subsequent migration from USA (<0.5 kya) |
|---|---|---|
| Akaike weight | 0.503 | 0.493 |
| Age of migration from USA | 301 (upper limit 500) | 158 (upper limit 467) |
| Migration pulse probability | 57.50% | 57.82% |
| Divergence age between Cuba and southern Mexico populations | 855 (constrained 500-5,000) | 467 (upper limit 500) |
| Divergence age between northern and southern Mexico populations | 333,786 | 334,693 |
| Divergence age between USA and ancestral Mexico population | 333,786 | 334,693 |
| Divergence age between Black-throated and ancestral Northern Bobwhite population | 1,562,873 (upper limit 1.563 M) | 1,562,902 (upper limit 1.563 M) |
| Cuba $N_e$ | 752 | 393 |
| Northern Mexico $N_e$ | 1,290,940 | 1,310,580 |
| Southern Mexico $N_e$ | 1,108,260 | 1,098,556 |
| USA $N_e$ | 916,239 | 927,960 |
| Black-throated Bobwhite $N_e$ | 2,213,379 | 2,262,612 |

**Table S6.** Samples used for RADcap bait design**.** Cat. No. = Catalog Number / Accession Number. Samples listed in bold are also included in the present study. Subspecies identifications are taken from the collection where the specimen is accessioned and may have been assigned by locality without detailed diagnosis.

| Species | Subspecies/ Subpopulation | Museum | Cat. No. | Locality |
|---|---|---|---|---|
| *C. cristatus* | *sonnini* | YPM | 137007 | Sipaliwini District, Suriname |
| *C. cristatus* | *sonnini* | YPM | 137076 | Sipaliwini District, Suriname |
| *C. cristatus* | *sonnini* | YPM | 137159 | Sipaliwini District, Suriname |
| *C. cristatus* | *sonnini* | YPM | 137267 | Sipaliwini District, Suriname |
| *C. cristatus* | *sonnini* | YPM | 137466 | Sipaliwini District, Suriname |
| *C. cristatus* | *sonnini* | YPM | 137592 | Sipaliwini District, Suriname |
| *C. cristatus* | *sonnini* | USNM | 610117 | Upper Demerara-Berbice Region, Guyana |
| *C. cristatus* | *sonnini* | USNM | 610185 | Upper Demerara-Berbice Region, Guyana |
| -*C. cristatus* | *sonnini* | USNM | 616592 | Upper Demerara-Berbice Region, Guyana |
| *C. cristatus* | *sonnini* | USNM | 621082 | Upper Demerara-Berbice Region, Guyana |
| *C. cristatus* | *sonnini* | USNM | 622136 | Upper Demerara-Berbice Region, Guyana |
| *C. cristatus* | *sonnini* | USNM | 622229 | Upper Demerara-Berbice Region, Guyana |
| *C. cristatus* | *sonnini* | USNM | 626076 | Upper Takutu-Upper Essequibo Region, Guyana |
| *C. cristatus* | *sonnini* | USNM | 632665 | Upper Takutu-Upper Essequibo Region, Guyana |
| *C. cristatus* | *sonnini* | USNM | 632839 | Upper Takutu-Upper Essequibo Region, Guyana |
| *C. [leucopogon] cristatus* | *hypoleucos* | USNM | 646829 | La Paz Department, El Salvador |
| *C. [leucopogon] cristatus* | *leucopogon* | UWBM | 103378 | Copán Department, Honduras |
| *C. nigrogularis* | *caboti* | KU | 89347 | Yucatán, Mexico |
| *C. virginianus* | *cubanensis* | LSU | 92069 | Abaco District, Bahamas |
| *C. virginianus* | *cubanensis* | LSU | 48983 | New Providence District, Bahamas |
| *C. virginianus* | *cubanensis* | LSU | 92443 | New Providence District, Bahamas |
| *C. virginianus* | *cubanensis* | LSU | 92487 | New Providence District, Bahamas |
| *C. virginianus* | *cubanensis* | LSU | 92488 | New Providence District, Bahamas |
| ***C. virginianus*** | ***floridanus*** | **LSU** | **48982** | **Florida, USA** |
| ***C. virginianus*** | ***mexicanus*** | **LSU** | **27329** | **Louisiana, USA** |
| ***C. virginianus*** | ***mexicanus*** | **LSU** | **3283** | **Louisiana, USA** |
| ***C. virginianus*** | ***mexicanus*** | **LSU** | **3713** | **Louisiana, USA** |
| ***C. virginianus*** | ***mexicanus*** | **LSU** | **64490** | **Louisiana, USA** |
| ***C. virginianus*** | ***mexicanus*** | **LSU** | **91876** | **Louisiana, USA** |
| ***C. virginianus*** | ***mexicanus*** | **LSU** | **91918** | **Louisiana, USA** |
| ***C. virginianus*** | ***mexicanus*** | **LSU** | **91919** | **Louisiana, USA** |
| ***C. virginianus*** | ***mexicanus*** | **LSU** | **91920** | **Louisiana, USA** |
| ***C. virginianus*** | ***nigripectus*** | **SDNHM** | **54282** | **Morelos, Mexico** |
| ***C. virginianus*** | ***nigripectus*** | **SDNHM** | **54283** | **Morelos, Mexico** |
| ***C. virginianus*** | ***nigripectus*** | **SDNHM** | **54284** | **Morelos, Mexico** |
| *C. virginianus* | *taylori* | KU | 21905 | Kansas, USA |
| *C. virginianus* | *taylori* | KU | 6517 | Kansas, USA |
| *C. virginianus* | *taylori* | KU | 7066 | Kansas, USA |
| *C. virginianus* | *taylori* | LSU | 62470 | Texas, USA |
| *C. virginianus* | *taylori* | LSU | 62471 | Texas, USA |
| *C. virginianus* | *texanus* | LSU | 36012 | Texas, USA |
| *C. virginianus* | *texanus* | LSU | 54855 | Texas, USA |
| *C. virginianus* | *texanus* | LSU | 57493 | Texas, USA |
| *C. virginianus* | *texanus* | LSU | 62466 | Texas, USA |
| *C. virginianus* | *virginianus* | LSU | 69174 | Florida, USA |
| *C. virginianus* | *virginianus* | LSU | 92369 | Florida, USA |

**Table S7.** Catalog numbers (from Table S1) of samples included in randomly-sampled SNAPP analyses. Five samples were selected randomly from each population using custom R code. Column headings refer to the trees shown in Figure 3. N Mex = Northern Mexico; S Mex = Southern Mexico.

| Population | Fig. 3A | Fig. 3B | Fig. 3C | Fig. 3D | Fig. 3E |
|---|---|---|---|---|---|
| Cuba | 280778 | 58011 | 280780 | 33315 | 114910 |
| Cuba | 114910 | 33311 | 33317 | 114912 | 280779 |
| Cuba | 33320 | 33317 | 33320 | 33313 | 280778 |
| Cuba | 33319 | 33313 | 114912 | 280779 | 33313 |
| Cuba | 114912 | 280779 | 33313 | 280778 | 58012 |
| N Mex | 7701 | 35576 | 53454 | 36726 | 36726 |
| N Mex | 53450 | 45554 | 40230 | 7700 | 45549 |
| N Mex | 45554 | 25606 | 25607 | 39389 | 39457 |
| N Mex | 57882 | 36726 | 7700 | 39457 | 7702 |
| N Mex | 7700 | 53453 | 53453 | 45549 | 36640 |
| S Mex | 17806 | 54283 | 46883 | 17806 | 17806 |
| S Mex | 2828 | 28759 | 4146 | 27130 | 3679 |
| S Mex | 8857 | 54282 | 48912 | 8857 | 48912 |
| S Mex | 46883 | 43826 | 8854 | 24445 | 3680 |
| S Mex | 43827 | 27130 | 20968 | 20968 | 54282 |
| USA | 48982 | 27329 | 270595 | 3283 | 54851 |
| USA | 91920 | 91919 | 270596 | 48982 | 270595 |
| USA | 3283 | 270594 | 69842 | 54851 | 48982 |
| USA | 69842 | 270595 | 54852 | 14154 | 27329 |
| USA | 3713 | 54852 | 270597 | 14153 | 54852 |

**Table S8.** Catalog numbers (from Table S1) of samples included in by-locality SNAPP analyses (Figure 4). All available samples from each Cuban population were included, and two samples from each mainland subspecies were randomly selected using custom R code. N Mex = Northern Mexico; S Mex = Southern Mexico.

| Population | Pinar del Río | Isla de la Juventud | Sancti Spíritus | Cienfuegos | Holguín |
|---|---|---|---|---|---|
| Cuba | 33315 | 33310 | 58010 | 280778 | 114910 |
| Cuba | 33316 | 33311 | 58011 | 280779 | 114911 |
| Cuba | 33317 | 33312 | 58012 | 280780 | 114912 |
| Cuba | 33319 | 33313 | - | 280781 | - |
| Cuba | 33320 | - | - | - | - |
| N Mex | 45554 | 45554 | 45549 | 45549 | 7701 |
| N Mex | 7700 | 7702 | 7701 | 7702 | 7702 |
| N Mex | 36726 | 25609 | 35576 | 35576 | 57882 |
| N Mex | 53453 | 53450 | 36726 | 39388 | 57884 |
| N Mex | 39437 | 39437 | 39437 | 39437 | 39437 |
| N Mex | 40200 | 40230 | 40231 | 40232 | 40200 |
| S Mex | 12490 | 12490 | 20996 | 20968 | 12490 |
| S Mex | 4145 | 20996 | 27065 | 4146 | 20968 |
| S Mex | 27131 | 28759 | 28759 | 27130 | 28759 |
| S Mex | 28759 | 61133 | 61133 | 28759 | 61133 |
| S Mex | 8857 | 17806 | 17807 | 17807 | 24445 |
| S Mex | 8858 | 8854 | 8858 | 3680 | 8854 |
| S Mex | 104507 | 104504 | 104507 | 43827 | 104504 |
| S Mex | 104508 | 104507 | 43826 | 44393 | 44393 |
| S Mex | 3679 | 3679 | 3679 | 54283 | 54282 |
| S Mex | 54282 | 54283 | 54284 | 54284 | 54283 |
| S Mex | 2828 | 35866 | 35866 | 46867 | 35866 |
| S Mex | 35867 | 35867 | 35867 | 46883 | 46883 |
| USA | 270594 | 270594 | 14156 | 14154 | 14156 |
| USA | 69842 | 270596 | 270595 | 14155 | 270596 |
| USA | 3713 | 41222 | 3713 | 64490 | 3283 |
| USA | 41222 | 91919 | 91918 | 91876 | 41222 |
| USA | 54852 | 54851 | 54851 | 54851 | 54852 |
| USA | 54854 | 54854 | 54852 | 54854 | 54854 |

**Table S9.** Parameters for 45 demographic models. All models had 11 parameters. For graphical depictions of each model, see Figure S2. N Mex = northern Mexico; S Mex = southern Mexico.

| Model No. | Model type | Founder Pop. | Founder timing constraints | Migrant Pop. | Migrant timing constraints |
|---|---|---|---|---|---|
| m01 | Single source | N Mex | <0.5 kya | N Mex | after founding event |
| m02 | Single source | N Mex | 0.5-5.0 kya | N Mex | after founding event |
| m03 | Single source | N Mex | 5.0-11.7 kya | N Mex | after founding event |
| m04 | Single source | N Mex | 11.7-23 kya | N Mex | after founding event |
| m05 | Single source | N Mex | 23.0-140.0 kya | N Mex | after founding event |
| m06 | Single source | S Mex | <0.5 kya | S Mex | after founding event |
| m07 | Single source | S Mex | 0.5-5.0 kya | S Mex | after founding event |
| m08 | Single source | S Mex | 5.0-11.7 kya | S Mex | after founding event |
| m09 | Single source | S Mex | 11.7-23 kya | S Mex | after founding event |
| m10 | Single source | S Mex | 23.0-140.0 kya | S Mex | after founding event |
| m11 | Single source | Mex | <0.5 kya | Mex | after founding event |
| m12 | Single source | Mex | 0.5-5.0 kya | Mex | after founding event |
| m13 | Single source | Mex | 5.0-11.7 kya | Mex | after founding event |
| m14 | Single source | Mex | 11.7-23 kya | Mex | after founding event |
| m15 | Single source | Mex | 23.0-140.0 kya | Mex | after founding event |
| m16 | Single source | USA | <0.5 kya | USA | after founding event |
| m17 | Single source | USA | 0.5-5.0 kya | USA | after founding event |
| m18 | Single source | USA | 5.0-11.7 kya | USA | after founding event |
| m19 | Single source | USA | 11.7-23 kya | USA | after founding event |
| m20 | Single source | USA | 23.0-140.0 kya | USA | after founding event |
| m21 | Multiple sources | N Mex | <0.5 kya | USA | <0.5 kya, after founding event |
| m22 | Multiple sources | N Mex | 0.5-5.0 kya | USA | <0.5 kya |
| m23 | Multiple sources | N Mex | 5.0-11.7 kya | USA | <0.5 kya |
| m24 | Multiple sources | N Mex | 11.7-23 kya | USA | <0.5 kya |
| m25 | Multiple sources | N Mex | 23.0-140.0 kya | USA | <0.5 kya |
| m26 | Multiple sources | S Mex | <0.5 kya | USA | <0.5 kya, after founding event |
| m27 | Multiple sources | S Mex | 0.5-5.0 kya | USA | <0.5 kya |
| m28 | Multiple sources | S Mex | 5.0-11.7 kya | USA | <0.5 kya |
| m29 | Multiple sources | S Mex | 11.7-23 kya | USA | <0.5 kya |
| m30 | Multiple sources | S Mex | 23.0-140.0 kya | USA | <0.5 kya |
| m31 | Multiple sources | Mex | <0.5 kya | USA | <0.5 kya, after founding event |
| m32 | Multiple sources | Mex | 0.5-5.0 kya | USA | <0.5 kya |
| m33 | Multiple sources | Mex | 5.0-11.7 kya | USA | <0.5 kya |
| m34 | Multiple sources | Mex | 11.7-23 kya | USA | <0.5 kya |
| m35 | Multiple sources | Mex | 23.0-140.0 kya | USA | <0.5 kya |
| m36 | Multiple sources | USA | <0.5 kya | N Mex | <0.5 kya, after founding event |
| m37 | Multiple sources | USA | 0.5-5.0 kya | N Mex | <0.5 kya |
| m38 | Multiple sources | USA | 5.0-11.7 kya | N Mex | <0.5 kya |
| m39 | Multiple sources | USA | 11.7-23 kya | N Mex | <0.5 kya |
| m40 | Multiple sources | USA | 23.0-140.0 kya | N Mex | <0.5 kya |
| m41 | Multiple sources | USA | <0.5 kya | S Mex | <0.5 kya, after founding event |
| m42 | Multiple sources | USA | 0.5-5.0 kya | S Mex | <0.5 kya |
| m43 | Multiple sources | USA | 5.0-11.7 kya | S Mex | <0.5 kya |
| m44 | Multiple sources | USA | 11.7-23 kya | S Mex | <0.5 kya |
| m45 | Multiple sources | USA | 23.0-140.0 kya | S Mex | <0.5 kya |

**Table S10.** Results of demographic model selection for ten initial runs (A-J). Likelihood scores, corrected AIC scores ($AIC_c$), difference between model $AIC_c$ and the lowest $AIC_c$ ($\Delta_i$), and Akaike weights ($w_i$) are given. All models have 11 parameters. Models are listed in descending order of Akaike weights; the confidence set of models for each run are listed in bold. For graphical depictions of each model, see Figure S2.

(**A) Results of Run 1**

| Model | Likelihood score | AICc | Δi | $w_i$ |
|---|---|---|---|---|
| **m27** | **-19242.91** | **38510.53** | **0.00** | **0.99** |
| m42 | -19247.96 | 38520.65 | 10.11 | 0.01 |
| m45 | -19318.77 | 38662.27 | 151.73 | 0.00 |
| m21 | -19351.64 | 38728.00 | 217.47 | 0.00 |
| m43 | -19353.87 | 38732.47 | 221.93 | 0.00 |
| m20 | -19364.45 | 38753.62 | 243.08 | 0.00 |
| m24 | -19377.22 | 38779.15 | 268.62 | 0.00 |
| m28 | -19384.72 | 38794.16 | 283.63 | 0.00 |
| m44 | -19401.29 | 38827.29 | 316.76 | 0.00 |
| m26 | -19409.95 | 38844.63 | 334.10 | 0.00 |
| m17 | -19451.49 | 38927.70 | 417.17 | 0.00 |
| m16 | -19458.73 | 38942.18 | 431.65 | 0.00 |
| m18 | -19470.74 | 38966.20 | 455.67 | 0.00 |
| m5 | -19474.87 | 38974.46 | 463.93 | 0.00 |
| m39 | -19526.12 | 39076.97 | 566.44 | 0.00 |
| m38 | -19539.37 | 39103.46 | 592.93 | 0.00 |
| m30 | -19585.26 | 39195.24 | 684.71 | 0.00 |
| m36 | -19618.07 | 39260.86 | 750.32 | 0.00 |
| m23 | -19629.28 | 39283.28 | 772.74 | 0.00 |
| m19 | -19657.85 | 39340.43 | 829.89 | 0.00 |
| m10 | -19680.01 | 39384.75 | 874.21 | 0.00 |
| m29 | -19696.99 | 39418.70 | 908.17 | 0.00 |
| m4 | -19698.77 | 39422.27 | 911.74 | 0.00 |
| m25 | -19706.28 | 39437.28 | 926.75 | 0.00 |
| m3 | -19736.23 | 39497.18 | 986.65 | 0.00 |
| m1 | -19747.82 | 39520.37 | 1009.83 | 0.00 |
| m40 | -19812.42 | 39649.56 | 1139.02 | 0.00 |
| m34 | -19880.60 | 39785.92 | 1275.39 | 0.00 |
| m41 | -19901.99 | 39828.70 | 1318.16 | 0.00 |
| m37 | -19936.88 | 39898.48 | 1387.95 | 0.00 |
| m15 | -19939.45 | 39903.62 | 1393.08 | 0.00 |
| m12 | -23851.07 | 47726.86 | 9216.32 | 0.00 |
| m14 | -20299.45 | 40623.63 | 2113.10 | 0.00 |
| m32 | -20041.26 | 40107.24 | 1596.71 | 0.00 |
| m7 | -20330.93 | 40686.58 | 2176.04 | 0.00 |
| m9 | -20146.21 | 40317.13 | 1806.60 | 0.00 |
| m11 | -20928.60 | 41881.92 | 3371.39 | 0.00 |
| m13 | -20305.83 | 40636.37 | 2125.84 | 0.00 |
| m22 | -20937.39 | 41899.51 | 3388.98 | 0.00 |
| m2 | -20217.45 | 40459.61 | 1949.08 | 0.00 |
| m31 | -20218.98 | 40462.67 | 1952.14 | 0.00 |
| m33 | -20302.17 | 40629.07 | 2118.54 | 0.00 |
| m35 | -20013.60 | 40051.93 | 1541.39 | 0.00 |
| m6 | -20037.22 | 40099.16 | 1588.63 | 0.00 |
| m8 | -20022.19 | 40069.09 | 1558.56 | 0.00 |

**(B) Results of Run 2**

| Model | Likelihood score | AICc | Δi | $w_i$ |
|---|---|---|---|---|
| **m41** | **-19243.60** | **38511.92** | **0.00** | **1.00** |
| m28 | -19304.76 | 38634.25 | 122.33 | 0.00 |
| m40 | -19342.25 | 38709.22 | 197.30 | 0.00 |
| m20 | -19348.35 | 38721.42 | 209.50 | 0.00 |
| m43 | -19350.26 | 38725.25 | 213.33 | 0.00 |
| m23 | -19350.54 | 38725.80 | 213.88 | 0.00 |
| m21 | -19351.33 | 38727.38 | 215.47 | 0.00 |
| m45 | -19355.07 | 38734.86 | 222.94 | 0.00 |
| m27 | -19363.47 | 38751.67 | 239.75 | 0.00 |
| m22 | -19374.43 | 38773.58 | 261.66 | 0.00 |
| m24 | -19404.44 | 38833.60 | 321.68 | 0.00 |
| m19 | -19414.22 | 38853.17 | 341.25 | 0.00 |
| m18 | -19445.89 | 38916.50 | 404.59 | 0.00 |
| m17 | -19447.89 | 38920.50 | 408.58 | 0.00 |
| m37 | -19448.65 | 38922.02 | 410.10 | 0.00 |
| m5 | -19474.88 | 38974.49 | 462.57 | 0.00 |
| m16 | -19478.73 | 38982.18 | 470.26 | 0.00 |
| m38 | -19502.11 | 39028.94 | 517.02 | 0.00 |
| m30 | -19566.31 | 39157.34 | 645.42 | 0.00 |
| m39 | -19573.71 | 39172.13 | 660.22 | 0.00 |
| m25 | -19602.12 | 39228.97 | 717.05 | 0.00 |
| m44 | -19625.57 | 39275.86 | 763.94 | 0.00 |
| m35 | -19669.38 | 39363.47 | 851.55 | 0.00 |
| m36 | -19678.19 | 39381.10 | 869.18 | 0.00 |
| m15 | -19688.54 | 39401.79 | 889.88 | 0.00 |
| m29 | -19713.74 | 39452.20 | 940.28 | 0.00 |
| m4 | -19724.12 | 39472.95 | 961.04 | 0.00 |
| m3 | -19737.16 | 39499.04 | 987.12 | 0.00 |
| m2 | -19742.82 | 39510.36 | 998.44 | 0.00 |
| m1 | -19747.85 | 39520.42 | 1008.50 | 0.00 |
| m34 | -19827.49 | 39679.71 | 1167.79 | 0.00 |
| m33 | -19897.64 | 39820.01 | 1308.09 | 0.00 |
| m10 | -19907.71 | 39840.14 | 1328.22 | 0.00 |
| m12 | -20835.91 | 41696.54 | 3184.62 | 0.00 |
| m14 | -20241.59 | 40507.89 | 1995.98 | 0.00 |
| m32 | -24582.50 | 49189.71 | 10677.79 | 0.00 |
| m7 | -20073.43 | 40171.59 | 1659.67 | 0.00 |
| m9 | -20125.68 | 40276.09 | 1764.17 | 0.00 |
| m11 | -20395.18 | 40815.07 | 2303.16 | 0.00 |
| m13 | -22448.86 | 44922.43 | 6410.51 | 0.00 |
| m26 | -20293.91 | 40612.54 | 2100.62 | 0.00 |
| m31 | -19954.19 | 39933.10 | 1421.18 | 0.00 |
| m42 | -20895.46 | 41815.63 | 3303.71 | 0.00 |
| m6 | -20279.27 | 40583.27 | 2071.35 | 0.00 |
| m8 | -20116.70 | 40258.12 | 1746.20 | 0.00 |

**(C) Results of Run 3**

| Model | Likelihood score | AICc | Δi | $w_i$ |
|---|---|---|---|---|
| **m27** | **-19244.51** | **38513.75** | **0.00** | **1.00** |
| m28 | -19276.45 | 38577.62 | 63.87 | 0.00 |
| m41 | -19302.48 | 38629.68 | 115.94 | 0.00 |
| m29 | -19304.89 | 38634.50 | 120.76 | 0.00 |
| m30 | -19310.09 | 38644.90 | 131.15 | 0.00 |
| m43 | -19357.87 | 38740.47 | 226.72 | 0.00 |
| m21 | -19358.73 | 38742.18 | 228.44 | 0.00 |
| m23 | -19382.74 | 38790.21 | 276.46 | 0.00 |
| m36 | -19383.84 | 38792.40 | 278.66 | 0.00 |
| m20 | -19444.99 | 38914.71 | 400.96 | 0.00 |
| m17 | -19446.67 | 38918.06 | 404.32 | 0.00 |
| m19 | -19451.71 | 38928.14 | 414.40 | 0.00 |
| m16 | -19455.48 | 38935.68 | 421.94 | 0.00 |
| m35 | -19472.81 | 38970.34 | 456.60 | 0.00 |
| m5 | -19474.89 | 38974.51 | 460.76 | 0.00 |
| m25 | -19512.33 | 39049.39 | 535.64 | 0.00 |
| m44 | -19522.77 | 39070.25 | 556.51 | 0.00 |
| m38 | -19556.37 | 39137.45 | 623.71 | 0.00 |
| m45 | -19559.51 | 39143.73 | 629.99 | 0.00 |
| m24 | -19598.69 | 39222.10 | 708.35 | 0.00 |
| m18 | -19620.72 | 39266.16 | 752.42 | 0.00 |
| m37 | -19687.55 | 39399.83 | 886.08 | 0.00 |
| m4 | -19699.07 | 39422.87 | 909.12 | 0.00 |
| m3 | -19724.03 | 39472.77 | 959.03 | 0.00 |
| m40 | -19729.21 | 39483.15 | 969.41 | 0.00 |
| m22 | -19739.35 | 39503.41 | 989.67 | 0.00 |
| m2 | -19742.79 | 39510.30 | 996.56 | 0.00 |
| m1 | -19747.21 | 39519.14 | 1005.39 | 0.00 |
| m15 | -19776.55 | 39577.83 | 1064.08 | 0.00 |
| m10 | -22303.54 | 44631.80 | 6118.06 | 0.00 |
| m12 | -21490.83 | 43006.39 | 4492.64 | 0.00 |
| m14 | -20272.22 | 40569.17 | 2055.42 | 0.00 |
| m32 | -20123.41 | 40271.55 | 1757.80 | 0.00 |
| m34 | -19974.81 | 39974.34 | 1460.60 | 0.00 |
| m7 | -22002.00 | 44028.72 | 5514.97 | 0.00 |
| m9 | -20454.23 | 40933.18 | 2419.44 | 0.00 |
| m11 | -20438.41 | 40901.54 | 2387.79 | 0.00 |
| m13 | -20856.69 | 41738.11 | 3224.36 | 0.00 |
| m26 | -20672.42 | 41369.57 | 2855.82 | 0.00 |
| m31 | -19988.14 | 40001.00 | 1487.25 | 0.00 |
| m33 | -20184.27 | 40393.27 | 1879.52 | 0.00 |
| m39 | -20125.90 | 40276.52 | 1762.77 | 0.00 |
| m42 | -22505.50 | 45035.73 | 6521.98 | 0.00 |
| m6 | -20063.80 | 40152.32 | 1638.57 | 0.00 |
| m8 | -20136.61 | 40297.94 | 1784.19 | 0.00 |

**(D) Results of Run 4**

| Model | Likelihood score | AICc | Δi | $w_i$ |
|---|---|---|---|---|
| **m41** | **-19258.04** | **38540.79** | **0.00** | **1.00** |
| m29 | -19285.73 | 38596.18 | 55.39 | 0.00 |
| m20 | -19323.46 | 38671.65 | 130.85 | 0.00 |
| m27 | -19351.35 | 38727.43 | 186.63 | 0.00 |
| m21 | -19351.55 | 38727.82 | 187.03 | 0.00 |
| m23 | -19390.39 | 38805.51 | 264.72 | 0.00 |
| m36 | -19391.13 | 38806.98 | 266.19 | 0.00 |
| m26 | -19413.57 | 38851.86 | 311.06 | 0.00 |
| m37 | -19417.29 | 38859.29 | 318.50 | 0.00 |
| m18 | -19451.88 | 38928.48 | 387.69 | 0.00 |
| m17 | -19454.87 | 38934.46 | 393.67 | 0.00 |
| m5 | -19474.87 | 38974.47 | 433.67 | 0.00 |
| m44 | -19510.05 | 39044.83 | 504.04 | 0.00 |
| m24 | -19514.96 | 39054.65 | 513.86 | 0.00 |
| m25 | -19539.88 | 39104.48 | 563.68 | 0.00 |
| m45 | -19539.97 | 39104.66 | 563.87 | 0.00 |
| m39 | -19552.27 | 39129.26 | 588.47 | 0.00 |
| m42 | -19559.33 | 39143.38 | 602.59 | 0.00 |
| m28 | -19580.28 | 39185.28 | 644.49 | 0.00 |
| m10 | -19635.54 | 39295.79 | 755.00 | 0.00 |
| m16 | -19635.97 | 39296.66 | 755.87 | 0.00 |
| m19 | -19639.36 | 39303.44 | 762.65 | 0.00 |
| m40 | -19646.09 | 39316.90 | 776.10 | 0.00 |
| m15 | -19683.91 | 39392.55 | 851.76 | 0.00 |
| m3 | -19724.84 | 39474.40 | 933.61 | 0.00 |
| m4 | -19729.20 | 39483.12 | 942.33 | 0.00 |
| m2 | -19747.15 | 39519.02 | 978.23 | 0.00 |
| m1 | -19747.48 | 39519.69 | 978.90 | 0.00 |
| m33 | -19932.26 | 39889.25 | 1348.46 | 0.00 |
| m12 | -20429.72 | 40884.16 | 2343.37 | 0.00 |
| m14 | -20345.11 | 40714.94 | 2174.15 | 0.00 |
| m30 | -20311.05 | 40646.82 | 2106.03 | 0.00 |
| m32 | -20488.84 | 41002.40 | 2461.61 | 0.00 |
| m34 | -20809.35 | 41643.42 | 3102.63 | 0.00 |
| m38 | -20169.76 | 40364.25 | 1823.46 | 0.00 |
| m43 | -19998.50 | 40021.72 | 1480.93 | 0.00 |
| m7 | -29179.63 | 58383.98 | 19843.18 | 0.00 |
| m9 | -20019.76 | 40064.25 | 1523.45 | 0.00 |
| m11 | -22201.78 | 44428.29 | 5887.50 | 0.00 |
| m13 | -20363.63 | 40751.99 | 2211.19 | 0.00 |
| m22 | -27463.12 | 54950.97 | 16410.18 | 0.00 |
| m31 | -20050.84 | 40126.40 | 1585.61 | 0.00 |
| m35 | -20005.79 | 40036.30 | 1495.51 | 0.00 |
| m6 | -24191.95 | 48408.63 | 9867.84 | 0.00 |
| m8 | -20014.17 | 40053.06 | 1512.27 | 0.00 |

**(E) Results of Run 5**

| Model | Likelihood score | AICc | Δi | $w_i$ |
|---|---|---|---|---|
| **m26** | **-19249.97** | **38524.66** | **0.00** | **1.00** |
| m20 | -19312.71 | 38650.15 | 125.49 | 0.00 |
| m40 | -19340.74 | 38706.21 | 181.55 | 0.00 |
| m36 | -19351.27 | 38727.26 | 202.60 | 0.00 |
| m42 | -19400.36 | 38825.45 | 300.79 | 0.00 |
| m44 | -19420.77 | 38866.27 | 341.60 | 0.00 |
| m18 | -19431.87 | 38888.45 | 363.79 | 0.00 |
| m35 | -19461.02 | 38946.77 | 422.11 | 0.00 |
| m16 | -19462.50 | 38949.72 | 425.06 | 0.00 |
| m38 | -19473.21 | 38971.14 | 446.48 | 0.00 |
| m21 | -19473.35 | 38971.42 | 446.76 | 0.00 |
| m37 | -19516.73 | 39058.19 | 533.53 | 0.00 |
| m29 | -19533.80 | 39092.32 | 567.66 | 0.00 |
| m19 | -19534.26 | 39093.24 | 568.57 | 0.00 |
| m41 | -19540.07 | 39104.86 | 580.19 | 0.00 |
| m39 | -19582.44 | 39189.61 | 664.95 | 0.00 |
| m5 | -19583.77 | 39192.25 | 667.59 | 0.00 |
| m17 | -19584.75 | 39194.22 | 669.56 | 0.00 |
| m27 | -19588.27 | 39201.27 | 676.60 | 0.00 |
| m23 | -19588.51 | 39201.73 | 677.07 | 0.00 |
| m24 | -19599.46 | 39223.64 | 698.97 | 0.00 |
| m45 | -19599.51 | 39223.75 | 699.09 | 0.00 |
| m30 | -19637.08 | 39298.89 | 774.23 | 0.00 |
| m22 | -19641.04 | 39306.80 | 782.14 | 0.00 |
| m43 | -19684.48 | 39393.68 | 869.02 | 0.00 |
| m4 | -19704.21 | 39433.15 | 908.48 | 0.00 |
| m3 | -19724.02 | 39472.76 | 948.10 | 0.00 |
| m2 | -19752.50 | 39529.72 | 1005.05 | 0.00 |
| m15 | -19757.85 | 39540.43 | 1015.77 | 0.00 |
| m25 | -19827.39 | 39679.50 | 1154.84 | 0.00 |
| m1 | -19902.58 | 39829.88 | 1305.22 | 0.00 |
| m32 | -19939.04 | 39902.80 | 1378.14 | 0.00 |
| m33 | -19950.79 | 39926.29 | 1401.63 | 0.00 |
| m10 | -20012.41 | 40049.55 | 1524.89 | 0.00 |
| m12 | -20441.59 | 40907.89 | 2383.23 | 0.00 |
| m14 | -20799.47 | 41623.66 | 3099.00 | 0.00 |
| m34 | -19978.47 | 39981.67 | 1457.00 | 0.00 |
| m7 | -26128.40 | 52281.52 | 13756.85 | 0.00 |
| m9 | -21148.42 | 42321.55 | 3796.89 | 0.00 |
| m11 | -25586.94 | 51198.60 | 12673.94 | 0.00 |
| m13 | -20310.23 | 40645.19 | 2120.53 | 0.00 |
| m28 | -20205.19 | 40435.10 | 1910.44 | 0.00 |
| m31 | -19966.51 | 39957.74 | 1433.08 | 0.00 |
| m6 | -20279.46 | 40583.64 | 2058.97 | 0.00 |
| m8 | -20449.90 | 40924.53 | 2399.86 | 0.00 |

**(F) Results of Run 6**

| Model | Likelihood score | AICc | Δi | $w_i$ |
|---|---|---|---|---|
| **m27** | **-19238.65** | **38502.02** | **0.00** | **1.00** |
| m42 | -19246.85 | 38518.42 | 16.40 | 0.00 |
| m28 | -19263.31 | 38551.35 | 49.33 | 0.00 |
| m29 | -19275.06 | 38574.84 | 72.82 | 0.00 |
| m20 | -19316.75 | 38658.22 | 156.21 | 0.00 |
| m37 | -19372.79 | 38770.30 | 268.28 | 0.00 |
| m41 | -19391.74 | 38808.19 | 306.18 | 0.00 |
| m24 | -19394.16 | 38813.04 | 311.02 | 0.00 |
| m21 | -19397.54 | 38819.80 | 317.78 | 0.00 |
| m19 | -19414.89 | 38854.50 | 352.48 | 0.00 |
| m22 | -19435.06 | 38894.83 | 392.81 | 0.00 |
| m18 | -19436.47 | 38897.65 | 395.64 | 0.00 |
| m30 | -19443.42 | 38911.56 | 409.55 | 0.00 |
| m16 | -19462.38 | 38949.47 | 447.46 | 0.00 |
| m5 | -19474.89 | 38974.51 | 472.49 | 0.00 |
| m39 | -19488.33 | 39001.39 | 499.37 | 0.00 |
| m17 | -19488.80 | 39002.31 | 500.30 | 0.00 |
| m26 | -19509.60 | 39043.92 | 541.90 | 0.00 |
| m38 | -19532.06 | 39088.84 | 586.83 | 0.00 |
| m40 | -19532.19 | 39089.11 | 587.09 | 0.00 |
| m43 | -19537.74 | 39100.20 | 598.18 | 0.00 |
| m25 | -19592.34 | 39209.39 | 707.37 | 0.00 |
| m36 | -19609.04 | 39242.81 | 740.79 | 0.00 |
| m4 | -19698.86 | 39422.44 | 920.42 | 0.00 |
| m23 | -19725.31 | 39475.34 | 973.32 | 0.00 |
| m3 | -19726.65 | 39478.02 | 976.00 | 0.00 |
| m2 | -19742.55 | 39509.81 | 1007.79 | 0.00 |
| m1 | -19748.96 | 39522.65 | 1020.63 | 0.00 |
| m44 | -19822.71 | 39670.13 | 1168.11 | 0.00 |
| m15 | -19851.17 | 39727.06 | 1225.04 | 0.00 |
| m31 | -19928.66 | 39882.05 | 1380.03 | 0.00 |
| m33 | -19946.61 | 39917.95 | 1415.93 | 0.00 |
| m10 | -19989.81 | 40004.34 | 1502.32 | 0.00 |
| m12 | -20430.63 | 40885.97 | 2383.96 | 0.00 |
| m14 | -20213.30 | 40451.32 | 1949.30 | 0.00 |
| m32 | -20600.67 | 41226.06 | 2724.05 | 0.00 |
| m34 | -20159.26 | 40343.24 | 1841.22 | 0.00 |
| m45 | -20144.91 | 40314.55 | 1812.53 | 0.00 |
| m7 | -20124.16 | 40273.05 | 1771.03 | 0.00 |
| m9 | -22514.71 | 45054.14 | 6552.12 | 0.00 |
| m11 | -20433.36 | 40891.43 | 2389.42 | 0.00 |
| m13 | -20334.81 | 40694.34 | 2192.33 | 0.00 |
| m35 | -20018.61 | 40061.95 | 1559.93 | 0.00 |
| m6 | -20032.54 | 40089.81 | 1587.79 | 0.00 |
| m8 | -19998.33 | 40021.38 | 1519.36 | 0.00 |

#### (G) Results of Run 7

| Model | Likelihood score | AICc | Δi | $w_i$ |
|---|---|---|---|---|
| **m27** | **-19238.20** | **38501.13** | **0.00** | **1.00** |
| m41 | -19295.79 | 38616.30 | 115.17 | 0.00 |
| m29 | -19316.01 | 38656.74 | 155.61 | 0.00 |
| m20 | -19328.37 | 38681.46 | 180.33 | 0.00 |
| m45 | -19340.86 | 38706.43 | 205.30 | 0.00 |
| m23 | -19350.57 | 38725.86 | 224.73 | 0.00 |
| m36 | -19353.22 | 38731.17 | 230.04 | 0.00 |
| m22 | -19357.06 | 38738.84 | 237.71 | 0.00 |
| m40 | -19358.89 | 38742.51 | 241.38 | 0.00 |
| m42 | -19379.57 | 38783.86 | 282.73 | 0.00 |
| m18 | -19433.87 | 38892.47 | 391.34 | 0.00 |
| m19 | -19434.23 | 38893.18 | 392.05 | 0.00 |
| m26 | -19448.06 | 38920.84 | 419.71 | 0.00 |
| m43 | -19450.34 | 38925.39 | 424.27 | 0.00 |
| m16 | -19470.43 | 38965.58 | 464.45 | 0.00 |
| m25 | -19473.06 | 38970.84 | 469.71 | 0.00 |
| m5 | -19474.87 | 38974.47 | 473.34 | 0.00 |
| m17 | -19481.74 | 38988.20 | 487.07 | 0.00 |
| m38 | -19502.09 | 39028.90 | 527.77 | 0.00 |
| m30 | -19512.00 | 39048.73 | 547.60 | 0.00 |
| m35 | -19537.60 | 39099.93 | 598.80 | 0.00 |
| m37 | -19555.01 | 39134.74 | 633.61 | 0.00 |
| m21 | -19555.21 | 39135.14 | 634.01 | 0.00 |
| m28 | -19562.69 | 39150.11 | 648.98 | 0.00 |
| m44 | -19566.70 | 39158.13 | 657.00 | 0.00 |
| m24 | -19591.11 | 39206.94 | 705.81 | 0.00 |
| m39 | -19664.14 | 39353.00 | 851.87 | 0.00 |
| m4 | -19722.40 | 39469.53 | 968.40 | 0.00 |
| m2 | -19742.49 | 39509.69 | 1008.57 | 0.00 |
| m3 | -19744.73 | 39514.19 | 1013.06 | 0.00 |
| m1 | -19748.73 | 39522.19 | 1021.06 | 0.00 |
| m15 | -19805.50 | 39635.71 | 1134.59 | 0.00 |
| m33 | -19925.67 | 39876.07 | 1374.94 | 0.00 |
| m32 | -19927.54 | 39879.81 | 1378.68 | 0.00 |
| m10 | -20655.83 | 41336.37 | 2835.24 | 0.00 |
| m12 | -20383.56 | 40791.84 | 2290.71 | 0.00 |
| m14 | -20080.92 | 40186.57 | 1685.44 | 0.00 |
| m34 | -20012.95 | 40050.63 | 1549.50 | 0.00 |
| m7 | -19989.97 | 40004.67 | 1503.54 | 0.00 |
| m9 | -20262.70 | 40550.13 | 2049.00 | 0.00 |
| m11 | -20564.57 | 41153.87 | 2652.74 | 0.00 |
| m13 | -20687.42 | 41399.55 | 2898.42 | 0.00 |
| m31 | -19949.23 | 39923.18 | 1422.05 | 0.00 |
| m6 | -19993.54 | 40011.80 | 1510.67 | 0.00 |
| m8 | -20509.91 | 41044.53 | 2543.41 | 0.00 |

#### (H) Results of Run 8

| Model | Likelihood score | AICc | Δi | $w_i$ |
|---|---|---|---|---|
| **m26** | **-19271.86** | **38568.44** | **0.00** | **1.00** |
| m29 | -19310.54 | 38645.80 | 77.36 | 0.00 |
| m23 | -19350.97 | 38726.66 | 158.21 | 0.00 |
| m43 | -19366.82 | 38758.37 | 189.93 | 0.00 |
| m20 | -19379.25 | 38783.22 | 214.78 | 0.00 |
| m37 | -19383.46 | 38791.65 | 223.20 | 0.00 |
| m41 | -19386.25 | 38797.22 | 228.78 | 0.00 |
| m40 | -19394.30 | 38813.32 | 244.88 | 0.00 |
| m24 | -19408.10 | 38840.92 | 272.48 | 0.00 |
| m21 | -19410.90 | 38846.53 | 278.09 | 0.00 |
| m27 | -19424.21 | 38873.14 | 304.70 | 0.00 |
| m35 | -19425.75 | 38876.23 | 307.79 | 0.00 |
| m17 | -19451.93 | 38928.57 | 360.13 | 0.00 |
| m16 | -19462.51 | 38949.74 | 381.30 | 0.00 |
| m42 | -19475.92 | 38976.57 | 408.13 | 0.00 |
| m45 | -19486.78 | 38998.28 | 429.84 | 0.00 |
| m39 | -19523.72 | 39072.16 | 503.72 | 0.00 |
| m25 | -19532.45 | 39089.62 | 521.17 | 0.00 |
| m44 | -19534.01 | 39092.74 | 524.30 | 0.00 |
| m36 | -19588.19 | 39201.10 | 632.66 | 0.00 |
| m22 | -19634.46 | 39293.64 | 725.20 | 0.00 |
| m19 | -19639.60 | 39303.92 | 735.48 | 0.00 |
| m30 | -19651.31 | 39327.35 | 758.91 | 0.00 |
| m18 | -19697.84 | 39420.40 | 851.96 | 0.00 |
| m4 | -19698.61 | 39421.94 | 853.50 | 0.00 |
| m3 | -19726.99 | 39478.70 | 910.26 | 0.00 |
| m15 | -19740.72 | 39506.15 | 937.71 | 0.00 |
| m1 | -19751.67 | 39528.06 | 959.62 | 0.00 |
| m28 | -19940.96 | 39906.64 | 1338.20 | 0.00 |
| m10 | -22222.25 | 44469.22 | 5900.78 | 0.00 |
| m12 | -20368.95 | 40762.62 | 2194.18 | 0.00 |
| m14 | -20387.70 | 40800.13 | 2231.69 | 0.00 |
| m32 | -21044.67 | 42114.07 | 3545.63 | 0.00 |
| m34 | -20018.00 | 40060.73 | 1492.29 | 0.00 |
| m38 | -20043.80 | 40112.32 | 1543.88 | 0.00 |
| m5 | -24634.35 | 49293.41 | 10724.97 | 0.00 |
| m7 | -20283.30 | 40591.32 | 2022.87 | 0.00 |
| m9 | -20117.83 | 40260.39 | 1691.95 | 0.00 |
| m11 | -22962.03 | 45948.78 | 7380.34 | 0.00 |
| m13 | -20293.02 | 40610.77 | 2042.33 | 0.00 |
| m2 | -20021.54 | 40067.79 | 1499.35 | 0.00 |
| m31 | -20010.27 | 40045.25 | 1476.81 | 0.00 |
| m33 | -19981.56 | 39987.84 | 1419.40 | 0.00 |
| m6 | -20248.37 | 40521.45 | 1953.01 | 0.00 |
| m8 | -20129.59 | 40283.91 | 1715.47 | 0.00 |

**(I) Results of Run 9**

| Model | Likelihood score | AICc | Δi | $w_i$ |
|---|---|---|---|---|
| m26 | -19247.97 | 38520.66 | 0.00 | 1.00 |
| m28 | -19278.08 | 38580.88 | 60.21 | 0.00 |
| m41 | -19283.15 | 38591.03 | 70.36 | 0.00 |
| m20 | -19312.65 | 38650.02 | 129.36 | 0.00 |
| m43 | -19357.09 | 38738.89 | 218.23 | 0.00 |
| m24 | -19359.57 | 38743.86 | 223.20 | 0.00 |
| m36 | -19395.97 | 38816.66 | 295.99 | 0.00 |
| m22 | -19425.09 | 38874.89 | 354.23 | 0.00 |
| m19 | -19427.85 | 38880.41 | 359.75 | 0.00 |
| m21 | -19430.65 | 38886.02 | 365.36 | 0.00 |
| m39 | -19430.82 | 38886.37 | 365.70 | 0.00 |
| m44 | -19431.20 | 38887.12 | 366.45 | 0.00 |
| m18 | -19447.20 | 38919.12 | 398.46 | 0.00 |
| m17 | -19455.79 | 38936.31 | 415.64 | 0.00 |
| m38 | -19478.44 | 38981.61 | 460.94 | 0.00 |
| m16 | -19490.32 | 39005.36 | 484.69 | 0.00 |
| m25 | -19501.56 | 39027.84 | 507.18 | 0.00 |
| m30 | -19517.81 | 39060.35 | 539.69 | 0.00 |
| m10 | -19534.08 | 39092.88 | 572.21 | 0.00 |
| m45 | -19571.59 | 39167.90 | 647.24 | 0.00 |
| m42 | -19581.59 | 39187.90 | 667.24 | 0.00 |
| m5 | -19589.64 | 39204.01 | 683.34 | 0.00 |
| m35 | -19598.78 | 39222.28 | 701.62 | 0.00 |
| m40 | -19686.41 | 39397.54 | 876.88 | 0.00 |
| m23 | -19698.02 | 39420.75 | 900.09 | 0.00 |
| m4 | -19699.29 | 39423.30 | 902.64 | 0.00 |
| m29 | -19722.75 | 39470.22 | 949.55 | 0.00 |
| m3 | -19734.60 | 39493.92 | 973.26 | 0.00 |
| m37 | -19741.73 | 39508.18 | 987.51 | 0.00 |
| m2 | -19744.71 | 39514.15 | 993.48 | 0.00 |
| m1 | -19749.22 | 39523.17 | 1002.50 | 0.00 |
| m31 | -19928.63 | 39881.98 | 1361.32 | 0.00 |
| m12 | -21439.93 | 42904.58 | 4383.92 | 0.00 |
| m14 | -20186.27 | 40397.25 | 1876.59 | 0.00 |
| m27 | -29854.37 | 59733.46 | 21212.79 | 0.00 |
| m32 | -20112.85 | 40250.41 | 1729.75 | 0.00 |
| m34 | -20235.75 | 40496.22 | 1975.55 | 0.00 |
| m7 | -20074.25 | 40173.21 | 1652.55 | 0.00 |
| m9 | -20104.61 | 40233.95 | 1713.29 | 0.00 |
| m11 | -20419.92 | 40864.55 | 2343.89 | 0.00 |
| m13 | -20586.78 | 41198.27 | 2677.61 | 0.00 |
| m15 | -20291.23 | 40607.18 | 2086.51 | 0.00 |
| m33 | -20613.97 | 41252.67 | 2732.00 | 0.00 |
| m6 | -26420.64 | 52866.00 | 14345.33 | 0.00 |
| m8 | -20023.32 | 40071.36 | 1550.69 | 0.00 |

**(J) Results of Run 10**

| Model | Likelihood score | AICc | Δi | $w_i$ |
|---|---|---|---|---|
| m27 | -19238.69 | 38502.09 | 0.00 | 1.00 |
| m26 | -19255.85 | 38536.42 | 34.32 | 0.00 |
| m20 | -19313.76 | 38652.24 | 150.14 | 0.00 |
| m43 | -19365.83 | 38756.39 | 254.30 | 0.00 |
| m25 | -19400.47 | 38825.67 | 323.57 | 0.00 |
| m36 | -19418.37 | 38861.45 | 359.36 | 0.00 |
| m19 | -19424.09 | 38872.90 | 370.80 | 0.00 |
| m35 | -19431.17 | 38887.06 | 384.97 | 0.00 |
| m16 | -19461.02 | 38946.76 | 444.67 | 0.00 |
| m5 | -19474.90 | 38974.51 | 472.42 | 0.00 |
| m29 | -19485.04 | 38994.81 | 492.71 | 0.00 |
| m18 | -19490.61 | 39005.94 | 503.85 | 0.00 |
| m42 | -19493.96 | 39012.63 | 510.54 | 0.00 |
| m28 | -19526.20 | 39077.12 | 575.03 | 0.00 |
| m23 | -19550.14 | 39124.99 | 622.90 | 0.00 |
| m17 | -19562.47 | 39149.66 | 647.56 | 0.00 |
| m41 | -19570.49 | 39165.69 | 663.60 | 0.00 |
| m40 | -19572.47 | 39169.66 | 667.56 | 0.00 |
| m24 | -19577.03 | 39178.78 | 676.69 | 0.00 |
| m21 | -19590.06 | 39204.83 | 702.74 | 0.00 |
| m22 | -19592.49 | 39209.70 | 707.60 | 0.00 |
| m10 | -19607.34 | 39239.39 | 737.30 | 0.00 |
| m39 | -19616.20 | 39257.12 | 755.03 | 0.00 |
| m4 | -19698.50 | 39421.72 | 919.63 | 0.00 |
| m3 | -19738.59 | 39501.90 | 999.81 | 0.00 |
| m2 | -19765.14 | 39555.01 | 1052.92 | 0.00 |
| m1 | -19778.37 | 39581.46 | 1079.37 | 0.00 |
| m45 | -19788.66 | 39602.04 | 1099.94 | 0.00 |
| m30 | -19798.29 | 39621.30 | 1119.21 | 0.00 |
| m31 | -19926.32 | 39877.36 | 1375.27 | 0.00 |
| m12 | -29728.69 | 59482.11 | 20980.01 | 0.00 |
| m14 | -20189.60 | 40403.93 | 1901.83 | 0.00 |
| m32 | -20001.50 | 40027.72 | 1525.62 | 0.00 |
| m34 | -20181.75 | 40388.21 | 1886.12 | 0.00 |
| m38 | -20497.26 | 41019.24 | 2517.14 | 0.00 |
| m7 | -20190.83 | 40406.38 | 1904.28 | 0.00 |
| m9 | -19982.99 | 39990.69 | 1488.60 | 0.00 |
| m11 | -23540.55 | 47105.81 | 8603.72 | 0.00 |
| m13 | -20349.84 | 40724.40 | 2222.31 | 0.00 |
| m15 | -20061.66 | 40148.04 | 1645.94 | 0.00 |
| m33 | -20417.73 | 40860.19 | 2358.10 | 0.00 |
| m37 | -20096.88 | 40218.49 | 1716.39 | 0.00 |
| m44 | -20138.75 | 40302.22 | 1800.12 | 0.00 |
| m6 | -20936.28 | 41897.28 | 3395.18 | 0.00 |
| m8 | -20308.26 | 40641.24 | 2139.15 | 0.00 |

**Table S11.** Results of demographic model selection. Each model was run 100 times and the highest likelihood score for each was used for AIC model selection. The number of parameters (K), corrected AIC scores ($AIC_c$), difference between model $AIC_c$ and the lowest $AIC_c$ ($\Delta_i$), and Akaike weights ($w_i$) are given. Model numbers correspond to the graphical depictions in Figure S2.

| Model | Model description | K | Likelihood score | AICc | $\Delta_i$ | $w_i$ |
|---|---|---|---|---|---|---|
| m27 | Southern Mexico founder (0.5-5.0 kya), subsequent migration from U.S. (<0.5 kya) | 11 | -19238.203 | 38501.128 | 0.000 | 0.503 |
| m26 | Southern Mexico founder (<0.5 kya), subsequent migration from U.S. (<0.5 kya) | 11 | -19238.225 | 38501.171 | 0.043 | 0.493 |
| m41 | U.S. founder (<0.5 kya), subsequent migration from Southern Mexico (<0.5 kya) | 11 | -19243.078 | 38510.878 | 9.750 | 0.004 |

### SI References


1. D. Williford, *et al.*, Contemporary genetic structure of the northern bobwhite west of the Mississippi River. *The Journal of Wildlife Management* **78**, 914–929 (2014).
2. D. Williford, R. W. Deyoung, R. L. Honeycutt, L. A. Brennan, F. Hernández, Phylogeography of the bobwhite (Colinus) quails. *Wildlife Monographs* **193**, 1–49 (2016).
3. J. F. Salter, *et al.*, Historical specimens and the limits of subspecies phylogenomics in the New World quails (Odontophoridae). *Mol. Phylogenet. Evol.* **175**, 107559 (2022).
4. S. L. Hoffberg, *et al.*, RADcap: sequence capture of dual-digest RADseq libraries with identifiable duplicates and reduced missing data. *Mol. Ecol. Resour.* **16**, 1264–1278 (2016).
5. J. E. McCormack, W. L. E. Tsai, B. C. Faircloth, Sequence capture of ultraconserved elements from bird museum specimens. *Mol. Ecol. Resour.* (2015) https:/doi.org/10.1111/1755-0998.12466.
6. M. G. Harvey, B. T. Smith, T. C. Glenn, B. C. Faircloth, R. T. Brumfield, Sequence Capture versus Restriction Site Associated DNA Sequencing for Shallow Systematics. *Syst. Biol.* **65**, 910–924 (2016).
7. C. F. Graham, *et al.*, Impacts of degraded DNA on restriction enzyme associated DNA sequencing (RADSeq). *Mol. Ecol. Resour.* **15**, 1304–1315 (2015).
8. N. J. Bayona-Vásquez, *et al.*, Adapterama III: Quadruple-indexed, double/triple-enzyme RADseq libraries (2RAD/3RAD). *bioRxiv*, 205799 (2019).
9. N. A. Baird, *et al.*, Rapid SNP discovery and genetic mapping using sequenced RAD markers. *PLoS One* **3**, e3376 (2008).
10. N. Rohland, D. Reich, Cost-effective, high-throughput DNA sequencing libraries for multiplexed target capture. *Genome Res.* **22**, 939–946 (2012).
11. T. C. Glenn, *et al.*, Adapterama I: universal stubs and primers for 384 unique dual-indexed or 147,456 combinatorially-indexed Illumina libraries (iTru & iNext). *PeerJ* **7**, e7755 (2019).
12. B. C. Faircloth, "Running 3RAD Analysis" (2018) (November 1, 2018).
13. J. M. Catchen, A. Amores, P. Hohenlohe, W. Cresko, J. H. Postlethwait, Stacks: building and genotyping Loci de novo from short-read sequences. *G3* **1**, 171–182 (2011).
14. J. Catchen, P. A. Hohenlohe, S. Bassham, A. Amores, W. A. Cresko, Stacks: an analysis tool set for population genomics. *Mol. Ecol.* **22**, 3124–3140 (2013).
15. J. F. Salter, *et al.*, A Highly Contiguous Reference Genome for Northern Bobwhite (Colinus virginianus). *G3* **9**, 3929–3932 (2019).
16. H. Li, R. Durbin, Fast and accurate short read alignment with Burrows-Wheeler transform. *Bioinformatics* **25**, 1754–1760 (2009).
17. H. Li, *et al.*, The Sequence Alignment/Map format and SAMtools. *Bioinformatics* **25**, 2078–2079 (2009).
18. H. Li, A statistical framework for SNP calling, mutation discovery, association mapping and population genetical parameter estimation from sequencing data. *Bioinformatics* **27**, 2987–2993 (2011).
19. P. Danecek, *et al.*, The variant call format and VCFtools. *Bioinformatics* **27**, 2156–2158 (2011).
20. A. R. Quinlan, I. M. Hall, BEDTools: a flexible suite of utilities for comparing genomic features. *Bioinformatics* **26**, 841–842 (2010).
21. A. Moncrieff, *Thinning_function v1.0: a program to thin SNPs in VCF files* (2018) https:/doi.org/10.5281/zenodo.5498266.
22. J. Aerts, *et al.*, Extent of linkage disequilibrium in chicken. *Cytogenet. Genome Res.* **117**, 338–345 (2007).
23. A. F. A. Smit, R. Green, *RepeatMasker Open-4.0* (2013-2015).
24. M. Johnson, *et al.*, NCBI BLAST: a better web interface. *Nucleic Acids Res.* **36**, W5–9 (2008).
25. N. J. Bayona-Vásquez, *et al.*, Adapterama III: Quadruple-indexed, double/triple-enzyme RADseq libraries (2RAD/3RAD). *PeerJ* **7**, e7724 (2019).
26. T. C. Glenn, *et al.*, Adapterama I: Universal stubs and primers for thousands of dual-

indexed Illumina libraries (iTru & iNext). *bioRxiv*, 049114 (2016).
27. B. Bushnell, “BBMap: A fast, accurate, splice-aware aligner” (Lawrence Berkeley National Lab. (LBNL), Berkeley, CA (United States), 2014) (June 8, 2021).
28. B. C. Faircloth, Illumiprocessor: a trimmomatic wrapper for parallel adapter and quality trimming. *See http://dx. doi. org/10. 6079/J9ILL (accessed 4 November 2016)* (2013).
29. B. C. Faircloth, “Demultiplexing a Sequencing Run” (2018) (September 5, 2021).
30. J. F. Salter, B. C. Faircloth, “Running RADcap Analysis” (2021) (June 8, 2021).
31. G. A. Van der Auwera, B. D. O’Connor, Genomics in the Cloud: Using Docker, GATK, and WDL in Terra (2020).
32. J. F. Salter, B. C. Faircloth, “Running GATK in Parallel” (2021) (June 8, 2021).
33. A. McKenna, *et al.*, The Genome Analysis Toolkit: a MapReduce framework for analyzing next-generation DNA sequencing data. *Genome Res.* **20**, 1297–1303 (2010).
34. E. Frichot, F. Mathieu, T. Trouillon, G. Bouchard, O. François, Fast and efficient estimation of individual ancestry coefficients. *Genetics* **196**, 973–983 (2014).
35. J. Kamm, J. Terhorst, R. Durbin, Y. S. Song, Efficiently inferring the demographic history of many populations with allele count data. *J. Am. Stat. Assoc.* **115**, 1472–1487 (2020).
36. B. Chen, J. W. Cole, C. Grond-Ginsbach, Departure from Hardy Weinberg Equilibrium and Genotyping Error. *Front. Genet.* **8**, 167 (2017).
37. X. Yi, E. K. Latch, Nonrandom missing data can bias PCA inference of population genetic structure. *Mol. Ecol. Resour.* (2021) https:/doi.org/10.1111/1755-0998.13498.
38. B. Chattopadhyay, K. M. Garg, U. Ramakrishnan, Effect of diversity and missing data on genetic assignment with RAD-Seq markers. *BMC Res. Notes* **7**, 841 (2014).
39. E. Linck, C. J. Battey, Minor allele frequency thresholds strongly affect population structure inference with genomic data sets. *Mol. Ecol. Resour.* **19**, 639–647 (2019).
40. G. T. Marth, E. Czabarka, J. Murvai, S. T. Sherry, The allele frequency spectrum in genome-wide human variation data reveals signals of differential demographic history in three large world populations. *Genetics* **166**, 351–372 (2004).
41. B. J. Knaus, N. J. Grünwald, vcfr: a package to manipulate and visualize variant call format data in R. *Mol. Ecol. Resour.* **17**, 44–53 (2017).
42. T. Jombart, adegenet: a R package for the multivariate analysis of genetic markers. *Bioinformatics* **24**, 1403–1405 (2008).
43. R Core Team, R: A Language and Environment for Statistical Computing (2020).
44. T. Jombart, C. Collins, “A tutorial for Discriminant Analysis of Principal Components (DAPC) using adegenet 2.0.0” (2015) (September 5, 2021).
45. X. Zheng, *et al.*, A high-performance computing toolset for relatedness and principal component analysis of SNP data. *Bioinformatics* **28**, 3326–3328 (2012).
46. J. Goudet, HIERFSTAT, a package for to compute and test hierarchical F-statistics. *Mol. Ecol. Notes* **5**, 184–186 (2005).
47. G. Van Rossum, F. L. Drake, *Python 3 Reference Manual: (Python Documentation Manual Part 2)* (CreateSpace Independent Publishing Platform, 2009).
48. B. C. Faircloth, *private-alleles: Compute private alleles in one population relative to another* (Github, 2021) (September 7, 2021).
49. D. Bryant, R. Bouckaert, J. Felsenstein, N. A. Rosenberg, A. RoyChoudhury, Inferring species trees directly from biallelic genetic markers: bypassing gene trees in a full coalescent analysis. *Mol. Biol. Evol.* **29**, 1917–1932 (2012).
50. R. Bouckaert, *et al.*, BEAST 2: a software platform for Bayesian evolutionary analysis. *PLoS Comput. Biol.* **10**, e1003537 (2014).
51. S. K. Shakya, *vcf2SNAPP.R at master · shankarkshakya/mypackage* (Github, 2017) (September 5, 2021).
52. A. Rambaut, A. J. Drummond, D. Xie, G. Baele, M. A. Suchard, Posterior Summarization in Bayesian Phylogenetics Using Tracer 1.7. *Syst. Biol.* **67**, 901–904 (2018).
53. A. J. Drummond, S. Y. W. Ho, M. J. Phillips, A. Rambaut, Relaxed phylogenetics and dating with confidence. *PLoS Biol.* **4**, e88 (2006).
54. R. R. Bouckaert, DensiTree: making sense of sets of phylogenetic trees. *Bioinformatics* **26**, 1372–1373 (2010).
55. R. R. Bouckaert, J. Heled, DensiTree 2: Seeing Trees Through the Forest. *bioRxiv*,

012401 (2014).
56. J. Orihuela, An annotated list of Late Quaternary extinct birds of Cuba. *Ornitol. Neotrop.* **30**, 57–67 (2019).
57. J. C. Gundlach, *Contribucion á la ornitologia cubana,* (Imp. “La Antilla” de N. Cacho-Negrete, 1876).
58. F. M. Chapman, *Notes on Birds and Mammals Observed Near Trinidad, Cuba: With Remarks on the Origin of West Indian Bird-life* (order of the Trustees, American Museum of Natural History, 1892).
59. B. de las Casas, *Historia de las Indias* (Imprenta y litografia de I. Paz, 1877).
60. L. Allaire, “Archaeology of the Caribbean Region” in *The Cambridge History of the Native Peoples of the Americas*, F. Salomon, S. B. Schwartz, Eds. (Cambridge University Press, 1999).
61. K. M. Cohen, S. C. Finney, P. L. Gibbard, J.-X. Fan, The ICS international chronostratigraphic chart. *Episodes* **36**, 199–204 (2013).
62. F. Colleoni, C. Wekerle, J.-O. Näslund, J. Brandefelt, S. Masina, Constraint on the penultimate glacial maximum Northern Hemisphere ice topography (≈140 kyrs BP). *Quat. Sci. Rev.* **137**, 97–112 (2016).
63. Y. A. Halley, *et al.*, A draft de novo genome assembly for the northern bobwhite (Colinus virginianus) reveals evidence for a rapid decline in effective population size beginning in the Late Pleistocene. *PLoS One* **9**, e90240 (2014).
64. K. Nam, *et al.*, Molecular evolution of genes in avian genomes. *Genome Biol.* **11**, R68 (2010).
65. P. A. Hosner, E. L. Braun, R. T. Kimball, Land connectivity changes and global cooling shaped the colonization history and diversification of New World quail (Aves: Galliformes: Odontophoridae). *J. Biogeogr.* **42**, 1883–1895 (2015).
66. C. M. Hurvich, C.-L. Tsai, Regression and time series model selection in small samples. *Biometrika* **76**, 297–307 (1989).
67. C. M. Hurvich, C.-L. Tsai, A corrected Akaike information criterion for vector autoregressive model selection. *J. Time Ser. Anal.* **14**, 271–279 (1993).
68. K. P. Burnham, D. R. Anderson, A practical information-theoretic approach. *Model selection and multimodel inference* 2 (2002).
69. R. Royall, *Statistical Evidence: A Likelihood Paradigm* (CRC Press, 1997).
70. I. Gronau, M. J. Hubisz, B. Gulko, C. G. Danko, A. Siepel, Bayesian inference of ancient human demography from individual genome sequences. *Nat. Genet.* **43**, 1031–1034 (2011).
71. P. Danecek, *et al.*, Twelve years of SAMtools and BCFtools. *Gigascience* 10 (2021).
72. B. C. Faircloth, PHYLUCE is a software package for the analysis of conserved genomic loci. *Bioinformatics* **32**, 786–788 (2016).
73. A. H. Freedman, *et al.*, Genome sequencing highlights the dynamic early history of dogs. *PLoS Genet.* **10**, e1004016 (2014).
